# A distal promoter and aberrant splicing enable canonical translation of out-of-frame proteins in Huntington’s disease

**DOI:** 10.1101/2025.10.20.683480

**Authors:** Rachel Anderson, Tanishk Soni, Ankur Jain

## Abstract

Expanded CAG trinucleotide repeats cause more than a dozen neurodegenerative diseases, including Huntington’s disease (HD). In several disorders, these repeats are translated in multiple reading frames without identifiable AUG start codons. This process, called repeat-associated non-AUG (RAN) translation, generates aggregation-prone proteins, but its molecular basis remains unclear. Here, using affinity capture of CAG-repeat-containing RNAs, we identify a previously unannotated promoter ~33 kb upstream of the *HTT* gene. Transcripts initiating from this promoter undergo repeat-length-dependent aberrant splicing into exon 1 of the canonical *HTT*, embedding the CAG repeat in AUG-initiated frames encoding polyalanine and polyserine proteins. Comparative genomics indicates that this upstream promoter is primate-specific, helping explain inconsistencies across rodent models. Our findings establish a direct, AUG-dependent, splicing-mediated mechanism for out-of-frame repeat proteins in HD, expose critical gaps in current animal models, and identify novel splice junctions as potential therapeutic targets.

## Introduction

More than 50 degenerative disorders are caused by expansions of short tandem repeats^1^. Among these, an expanded CAG trinucleotide repeat is linked to at least 12 conditions, including dentatorubral-pallidoluysian atrophy, spinal and bulbar muscular atrophy, multiple spinocerebellar ataxias, and Huntington’s disease (HD)^1^. For each of these disorders, the length of the CAG repeat tract within the causative gene is polymorphic, and symptoms manifest only when the repeat number exceeds a critical threshold^1^. This threshold varies by disease, but longer repeats are typically associated with earlier onset and increased symptom severity^1–3^. In HD, the repeat lies within the coding sequence of the *Huntingtin* (*HTT*) transcript, where it encodes a polyglutamine tract. Proteins with expanded polyglutamine repeats are prone to aggregation, which is one proposed mechanism by which CAG repeats contribute to cellular toxicity^1–3^.

Beyond the canonical polyglutamine frame, expanded repeats also yield proteins in alternative reading frames without any apparent AUG initiation codons, in a process termed as repeat-associated non-AUG (RAN) translation^4–7^. RAN translation is observed across multiple repeat-expansion diseases, generating frame-specific, repeat-containing proteins. The mechanism by which these out of frame proteins arise is unclear. In HD, polyserine and polyalanine proteins are detectable in patient brain tissue and are proposed to contribute to neuronal toxicity^7,8^. Notably some mouse models of HD produce aberrant proteins^7,9^, while others do not^10,11^, suggesting that their production may depend on locus context rather than repeat length alone.

We recently showed that expanded CAG repeats can create *de novo* splice acceptor sites^12^. In appropriate sequence contexts, nearby donors can be spliced to the repeat, potentially embedding them within AUG-initiated open reading frames. This repeat-associated splicing followed by canonical translation can explain a subset of proteins previously attributed to RAN translation in plasmid-based minigene systems^12,13^. We therefore asked whether an analogous mechanism contributes to the production of out of frame peptides in HD. This notion presents an immediate challenge as the CAG tract is located only ~196 nt from the annotated 5’ end of the canonical *HTT* transcript, and this region lacks high-scoring splice donors (Supp. Fig. 1A). In principle, this hypothesis can be tested directly by examining full-length, repeat-containing transcripts to determine if aberrant splicing places the CAG repeats into unexpected, AUG-initiated open reading frames. The 5’ end of the *HTT* locus, however, is notoriously difficult to sequence^14^ due to its high GC-content and the presence of the repeat tract, resulting in sparse coverage in conventional RNA-seq datasets^15,16^. Moreover, any alternative transcripts driving this process may be rare, tissue-specific, or repeat-dependent. As a result, the true repertoire of repeat-containing transcripts produced in disease remains unclear.

To resolve this issue, we developed a method to enrich and sequence CAG repeat-containing transcripts, yielding >100-fold higher coverage across the *HTT* exon 1 region than standard workflows. Applying this method to patientderived cells led to an unexpected discovery: the *HTT* locus has an additional promoter located ~33 kb upstream from the canonical gene. Transcripts originating from this promoter undergo repeat-expansion-dependent mis-splicing. This aberrant splicing embeds the CAG repeat tract into sequence contexts that enable canonical, AUG-initiated translation of both polyalanine and polyserine proteins. These findings add to the growing body of evidence that expanded repeats disrupt RNA processing and provide a mechanistic explanation for aberrant, out-of-frame translation products in HD.

## Results

### Affinity purification enriches full-length RNAs with tandem CAG repeats

Standard RNA-sequencing approaches do not provide sufficient coverage of the *HTT* repeat region^14,17,18^. To overcome this limitation, we adapted the RNA affinity purification (RAP) procedure^19^ to enrich RNAs harboring expanded CAG tracts. In brief, we hybridized total RNA to a biotinylated bait oligonucleotide complementary to the repeat (6×CTG, Fig. 1A). Longer repeats permit binding of multiple bait oligonucleotides, and are expected to increase capture efficiency. We optimized the hybridization conditions so as to minimize non-specific binding and preserve RNA integrity (see Methods). Bound RNAs were captured on streptavidin-coated magnetic beads and thoroughly washed to remove non-specifically adsorbed RNAs and contaminating DNA (see Methods and Fig. 1A). A key modification in our protocol is in the elution step. Existing methods often elute the captured RNA with RNase H, which cleaves RNA in RNA:DNA duplexes and precludes recovery of full-length RNAs^19,20^. Instead, we eluted the RNA with 95% formamide. Formamide decreases the effective melting temperature of nucleic acid duplexes by about 0.5°C per percent^21^. Elution at 60°C allowed recovery of RNAs with long repeat tracts (up to 240×CAG; Supp. Fig. 1B) and RNA remained stable in these conditions^22^ (Supp. Fig. 1C).

**Figure 1.**
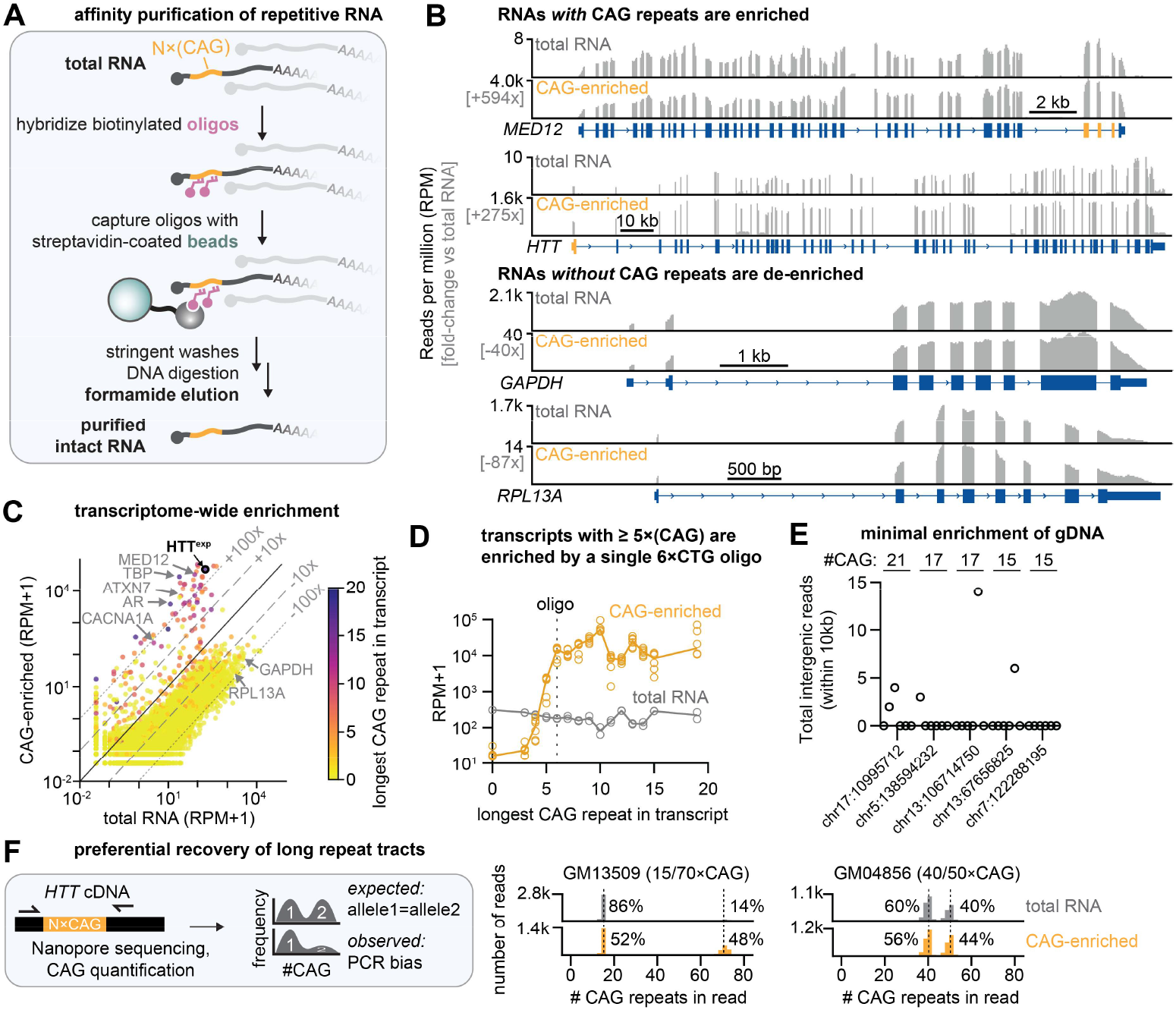
Affinity purification of tandem repeats enriches full-length RNAs with repetitive regions. **A**. Schematic for affinity purification of RNA with repeat tracts. Total RNA is hybridized to biotinylated bait oligonucleotide complementary to the repeat sequence. Next, bound RNAs are captured on streptavidin-coated magnetic beads and washed to remove weakly bound RNAs. Finally, RNA is eluted with a buffer containing 95% formamide, which disrupts nucleic acid duplexes and allows recovery of intact RNAs. **B**. Representative enrichment of RNAs with CAG repeats. Enriched RNA was subjected to short-read sequencing and aligned to hg38. The x-axis is the relative base position in the genome and the y-axis represents perbase read coverage. Gene schematics show the exons as rectangles separated by introns (thin lines). Exons containing ≥3×CAG repeats are highlighted in orange. **C**. Global enrichment of CAG repeats. Per-gene counts for total RNA sequenced from GM13509 (HD LCL) is plotted on the x-axis, and CAG-enriched GM13509 RNA is plotted on the y-axis. Each dot represents one gene, and is color coded to reflect the longest exonic CAG repeat in the gene. Dotted and dashed lines indicate 10 and 100-fold enrichment and depletion relative to total RNA input. Several genes associated with CAG repeat expansion disorders are annotated. **D**. Length-dependence of capture using a single 6×CTG bait. Reads per million mapped reads (RPM) abundance of RNAs with CAG repeats following short-read sequencing. Each dot represents the median read count of all transcripts annotated to carry a given number of CAG repeats in hg38 in a single sample, and the solid line shows the median across n=6 CAG-enriched and 2 total RNA inputs. The bait oligo used (corresponding to 6×CAG) is indicated as a dashed line. **E**. Total number of reads mapping within 10 kb on each side of intergenic CAG repeats, indicating minimal genomic DNA carryover. Each dot is one library. **F**. *Left:* Assay for determining repeat-specific enrichment. *HTT* cDNA from total RNA or CAG-enriched RNA was amplified using primers spanning the repeat, followed by long-read sequencing to a depth of >2000 reads. The short allele is over-represented due to PCR biases that favor amplification of shorter templates. *Right:* For two cell lines with known repeat lengths, the repeat length in each read was determined and plotted as a histogram. ~3.4-fold preferential enrichment of the long repeat in GM13509 was calculated as %pulldown_long_/%input_long_. Amplicon sequencing results are representative of 2 independent RNA isolations.

We first performed a proof-of-concept experiment on RNA from human lymphoblastoid cell lines (LCLs). Quantitative PCR revealed ~1,000-fold depletion of ribosomal RNAs and transcripts with ≤2×CAG repeats relative to input (normalizing by mass, Supp. Fig. 1D). In contrast, transcripts from genes with ≥6×CAG repeats were largely unchanged (~20% variation after normalization, Supp. Fig. 1D), indicating on-target enrichment. Small- and large-scale purifications (10 μg and 500-1500 μg input RNA, respectively) yielded comparable enrichment profiles (Supp. Fig. 1D). Recovery averaged around 0.02% of the input RNA by mass, consistent with the relative abundance of transcripts with expanded CAG repeats (~0.3% of RefSeq transcript annotations harbor ≥ 6×CAG repeats, Supp. Fig. 1E-F).

Short-read sequencing on CAG-enriched RNA confirmed selective enrichment without compromising data quality. In a representative sample, reads mapping to *MED12* and *HTT* increased by 594- and 275-fold, respectively relative to the total RNA (Fig. 1B). Similar results were observed for other CAG-repeat-containing transcripts (Fig. 1C, Supp. Fig. 1G). Read quality metrics, such as mismatch rate and the proportion of uniquely mapping reads were comparable to standard RNA-seq, while the proportion of reads containing CAG repeats increased by 4-fold (Supp. Fig. 1H-I). Transcripts with 5×CAG showed modest enrichment (median 9-fold), while those with ≥6×CAG showed strong enrichment (median 160-fold, Fig. 1C-D). In contrast, transcripts from genes lacking CAG tracts (≤2× tandem CAGs in exons, comprising ~99% of RefSeq transcript annotations) were depleted: *GAPDH* and *RPL13A* decreased by 40- and 87-fold, respectively (Fig. 1B-D). Similarly, highly abundant ribosomal and mitochondrial RNAs were depleted to a median of ~0.2% of mapped reads (Supp. Fig. 1I).

Intergenic regions with large genomic CAG repeats (≥15×CAG in hg38) were not detected, confirming RNA-specific capture rather than contaminating DNA (Fig. 1E, Supp. Fig. 1J). Gene-level counts after CAG enrichment were reproducible across six independently processed LCL cell lines (Pearson’s correlation coefficient between sample gene counts, r = 0.78-0.99, median = 0.86; Supp. Fig. 1K). Further preferential recovery of transcripts with long repeat tracts could be achieved by adjusting wash stringency, effectively depleting abundant mRNAs with short repeats (e.g. *SREBF2* with 6×CAG, see Methods and Supp. Fig. 1L). In a cell line with known *HTT* allele sizes (GM13509, 15/70×CAG), the expanded allele was enriched by 3.5-fold relative to the shorter allele (Fig. 1F). Our protocol also preserved RNA integrity, as evidenced by uniform sequencing coverage across the full gene body for enriched transcripts (Supp. Fig. 1M). Collectively, these data demonstrate that our protocol allows for robust and reproducible enrichment of full-length RNAs with tandem CAG repeats.

### The *HTT* gene has a distant upstream promoter

Applying this enrichment protocol to HD LCLs, we observed unexpected read coverage in the region between the *HTT* gene and its upstream neighbor *GRK4* extending ~33 kb upstream from the canonical *HTT* promoter (Fig. 2A). The coverage in this region is increased after CAG capture compared to total RNA, suggesting that these reads likely derive from CAG repeat-containing transcripts (Fig. 2A). This upstream coverage was specific to *HTT*, as such coverage was not detected at other genes with CAG repeats (Supp. Fig. 2A). *HTT* exon 1 harbors the only ≥5×CAG repeat tract within the entire *GRK4*-*HTT* locus (chr4:2,963,571-3,243,960 in hg38 coordinates), suggesting that these reads likely derive from the *HTT* CAG repeat.

**Figure 2.**
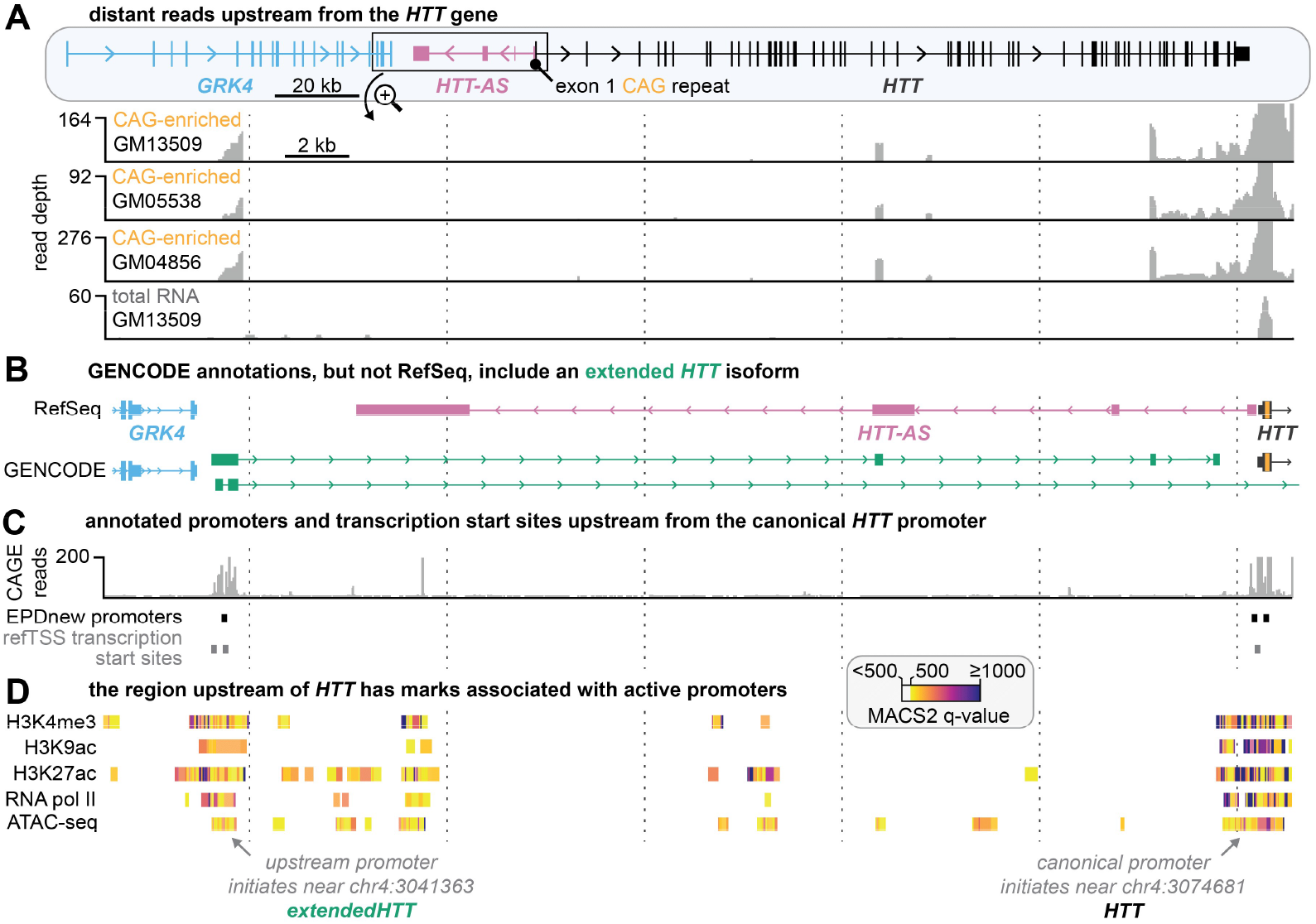
The HTT gene has an upstream promoter that generates extended HTT transcripts. **A**. *Top:* schematic for the *GRK4-HTT* region of chr4 (chr4:2963408-3244028). *Bottom:* CAG-enriched RNA-seq coverage from three HD LCLs for the zoomed-in region between *GRK4* and *HTT* indicated by a box (chr4:3037897-3075744). The Y-axis for each sample is normalized to 2% of the maximum read depth on *HTT* exon 1. Other panels in this figure use the same x-axis and vertical dashed lines are provided as visual aid. **B**. Gene annotations from NCBI RefSeq and GENCODE. The detected transcript is partially captured by GENCODE annotations, but not RefSeq. **C**. Cap Analysis of Gene Expression (CAGE) reads for the *GRK4*-*HTT* region support an upstream *HTT* promoter. CAGE reads indicate the 5’ ends of transcripts, often showing a distribution across a small region. Annotated upstream promoters (HTT_2 in EPDnew^27^) and transcription start sites (rfhg_55511.1, rfhg_55510.1 in RefTSS^28^) are indicated. **D**. ChIP-seq and accessibility data show promoter-like chromatin at the upstream site. Histone modifications associated with actively transcribed genes, RNA pol II occupancy, and chromatin accessibility were retrieved from ChIP-Atlas^29^ using a MACS (model-based analysis of ChIP-seq) q-value threshold for peak-calling significance of ≥500 (highly significant). The detected promoter region is indicated by arrows. Similar marks are present at the canonical *HTT* promoter.

We first excluded genomic DNA contamination as an explanation for these reads. Our protocol includes a DNase treatment step that should eliminate trace DNA contamination (see Methods). Consistent with this, intergenic regions with CAG repeats were depleted in our libraries: zero reads mapped within 10-kb of the five largest intergenic CAG repeats (Fig. 1E, Supp. Fig. 1J). Moreover, many upstream reads exhibited gapped alignments, suggesting that the intervening regions had been removed by RNA splicing (Supp. Fig. 2A, and further characterized in Figure 3).

**Figure 3.**
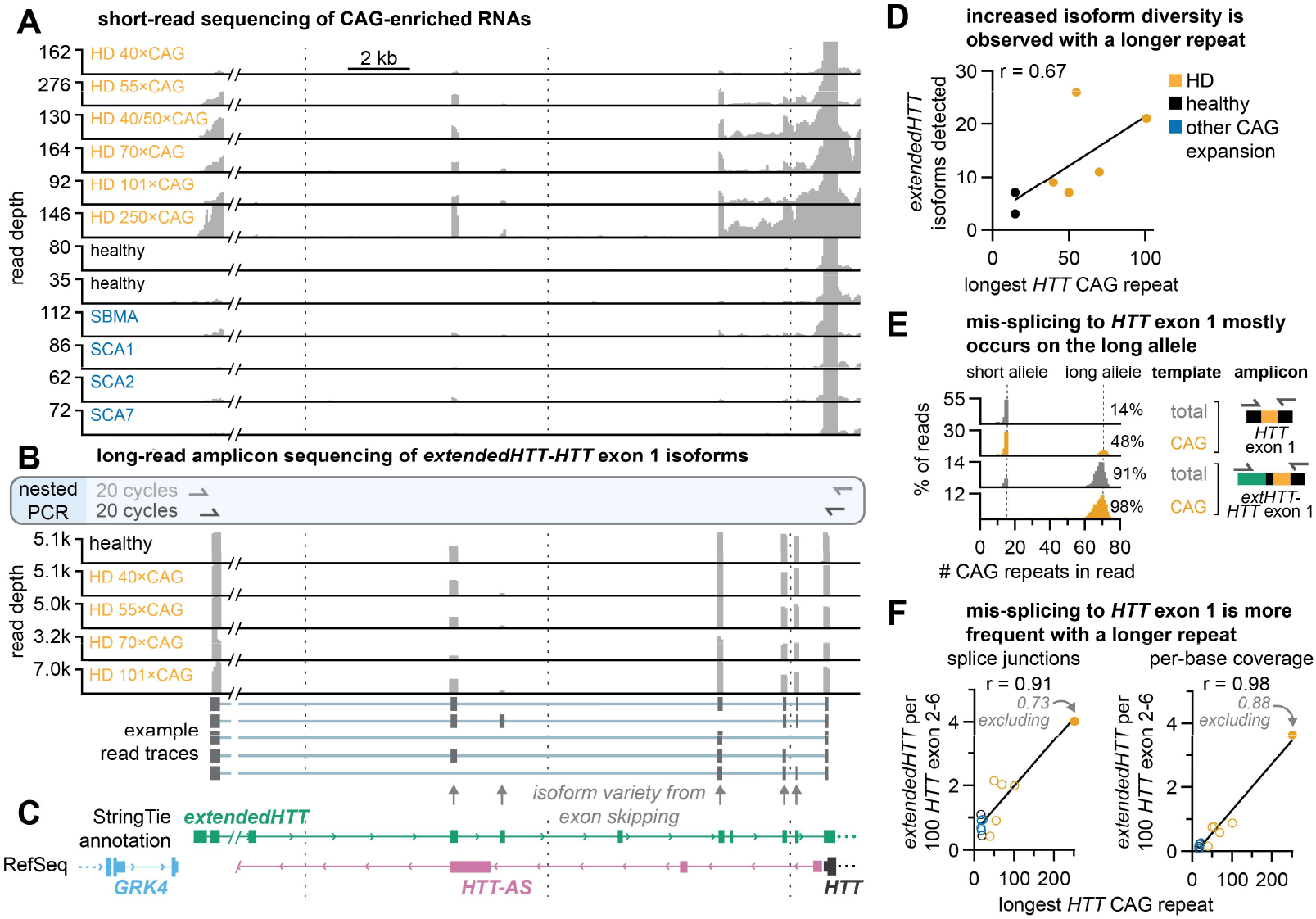
Expanded CAG repeats induce aberrant splicing of extendedHTT transcripts into HTT exon 1. **A**. CAG-enriched RNA sequencing for LCL cell lines. The x-axis shows genomic coordinates between *GRK4* and *HTT*, and is shared between panels B and C with vertical dashed lines provided as visual aid. The y-axis reflects read coverage per base, and was normalized to 2% of the maximum coverage on *HTT* exon 1 in that dataset. **B**. Oxford Nanopore long-read sequencing of cDNA amplicons prepared from CAG-enriched LCL RNA by two-step nested PCR. Not all exons are present in every transcript, as shown in sample reads, leading to isoform heterogeneity. **C**. Characterization of *extendedHTT* isoforms using StringTie. Aggregate CAG-enriched RNA-seq coverage for all samples from panels A and B were used for transcript assembly with StringTie. For simplicity, a single transcript isoform with all exons is indicated. **D**. Quantification of the number of isoforms detected (with a 10-read threshold) across long-read sequencing datasets. Legend is shared with panel F. **E**. Quantification of CAG repeat length in long-read sequencing of RT-PCR amplicons prepared from total RNA or CAG-enriched RNA. Amplicons targeting *HTT* exon 1 show that this method captures transcripts from both short and long alleles despite a PCR bias towards the short allele (as in Fig. 1F). However, most reads on the *extendedHTT-HTT* exon 1 isoform reflect the long allele. **F**. Quantification of *extendedHTT* abundance by frequency of splice junctions and per-base coverage of exons, relative to *HTT* exons 2-6, from CAG-enriched sequencing data. This normalization approach should account for increased recovery of transcripts with long repeats, as *extendedHTT* and *HTT* share a common CAG repeat.

We next sought to understand how transcripts potentially carrying both the *HTT* CAG tract and a far upstream region could be generated. Although the reads were in close proximity to *GRK4*, exons within the *GRK4* transcript were not enriched, ruling out transcriptional readthrough from the *GRK4* gene (Fig. 2A). During manual examination of the area with the genomic analysis tool IGV^23^, we noted that the regions of unexpected coverage did not overlap with any gene features in IGV’s default annotation schema (NCBI RefSeq, currently release 110^24^, Fig. 2A-B). Interestingly, however, recent GENCODE annotations include *HTT* transcript isoforms that overlap with the regions we detected (such as ENST00000680956.1, added in the 2019 v37 release of GENCODE^25^, see Methods and Fig. 2A-B). Although these isoforms have not been validated to the best of our knowledge, our CAG-enriched RNA-seq data is consistent with the GENCODE annotation and suggest that this transcript originates at roughly chr4:3041363 in hg38 coordinates, approximately 33kb upstream from the canonical *HTT* transcription start site (TSS; Fig. 2A-B, Supp. Fig. 2B).

Multiple independent lines of evidence support transcription initiation at this site. This site aligns with TSS tags from cap analysis of gene expression (CAGE) experiments probing the 5’ end of transcripts^26^, a promoter in the Eukaryotic Promoter Database^27^, and an annotated TSS in the refTSS database^28^ (Fig. 2C, Supp. Fig. 2B). Chromatin state analysis from ChIP-Atlas^29^ (a chromatin immunoprecipitation database) supports active transcription from this site, with the region near the putative TSS (chr4:3041363) showing enrichment in histone marks associated with promoters and actively transcribed genes, such as H3K4me3, H3K9ac, and H3K27ac^30^ (Fig. 2D). Last, this upstream region also displayed increased RNA polymerase II occupancy, consistent with promoter-proximal pausing, and increased chromatin accessibility, typical of transcriptionally active genes^31,32^ (Fig. 2D). These chromatin signatures closely resembled patterns observed at other genes with validated far-upstream promoters (Supp. Fig. 2C). Taken together, these orthogonal lines of evidence establish that there is an alternative promoter located ~33 kb upstream from the canonical *HTT* promoter. We designate transcripts originating from this promoter as *extendedHTT* isoforms to differentiate them from isoforms arising from the canonical *HTT* promoter (e.g., ENST00000355072.11).

### Expanded CAG repeats induce aberrant splicing of *extendedHTT* into *HTT* exon 1

The current GENCODE annotations indicate that upstream-initiated *HTT* isoforms (such as ENST00000680956.1) splice directly to exon 2, bypassing exon 1 and its CAG tract. Such exon 2 directed isoforms would not be captured in our CAG enrichment protocol. Our data, however, show robust enrichment of upstream-initiated transcripts in patient cells with numerous reads mapping across splice junctions that connect the upstream exons to canonical *HTT* exon 1 (Fig. 3A). These CAG-repeat containing *extendedHTT* transcripts were predominantly detected in HD carriers, but were minimally expressed in unaffected cells, indicative of repeat length dependent missplicing from upstream exons to the CAG repeat.

To determine whether these transcripts arise through canonical splicing mechanisms, we characterized their splice sites using complementary computational approaches. These transcripts contained up to ten exons upstream of canonical *HTT* (Fig. 3A). All putative splice junctions contained sequence motifs consistent with human 5’ and 3’ splice sites (‘GT’ and ‘AG’ dinucleotides, respectively, Supp. Fig. 3A, Supplemental Table 1). To confirm that these junctions represent functional splice sites, we applied MaxEntScan^33^, a splice-site prediction algorithm. Most of the identified splice sites in *extendedHTT* scored ≥ 0 (median 5’ MaxEnt score = 6.90 and 3’ score = 5.31, Supp. Fig. 3B, Supplemental Table 1), consistent with typical human splice sites (e.g., for 5’ sites, 97% score ≥ 0, median = 8.6, Supp. Fig. 3B). Additionally, the deep learning model SpliceAI^34^ assigned intermediate scores to many of these motifs, supporting the idea that they could participate in splicing (Supplemental Table 1).

In patient cells, the *extendedHTT* transcripts splice into the canonical *HTT* gene at exon 1 utilizing two main acceptor sites (Fig. 3A, Supp. Fig. 3C, Supplemental Table 1). The first acceptor is located within the 5’ untranslated region (UTR) of the canonical transcript (Supp. Fig. 3C). Interestingly, it overlaps with the SNP (single nucleotide polymorphism) rs146151652, where the minor allele modestly increases the splice site strength (C>T; 3’ MaxEnt score increases from 3.49 to 3.81). The second acceptor site is the CAG repeat itself (Supp. Fig. 3C-D), consistent with our prior report that tandem CAG repeats can act as splice acceptor sites^12^. Both acceptors position the *HTT* CAG repeat in an internal exon, explaining their enrichment by our CAG affinity purification protocol and indicating repeatdependent mis-splicing incorporates the repeat into *extendedHTT* transcripts.

To further characterize these transcripts, we generated RT-PCR amplicons using primers spanning from the 5’ end of *extendedHTT* to the 3’ end of the canonical *HTT* exon 1 (Fig. 3B) followed by Oxford Nanopore long-read sequencing. *De novo* transcript assembly using StringTie^35^, using a conservative threshold of ≥10 reads supporting a transcript, identified 38 distinct isoforms (see Supplemental Table 2) with variable usage across LCL lines (Fig. 3B-C). Strikingly, isoform diversity correlated with CAG repeat length and LCLs harboring longer repeats expressed more isoforms than those with shorter repeats (Pearson’s correlation coefficient, r, = 0.70, between CAG repeat number and the number of unique isoforms detected, Fig. 3D). Interestingly, our long-read sequencing data revealed that *extendedHTT* transcripts that are spliced to canonical *HTT* exon 1 almost exclusively originate from the expanded allele (Fig. 3E, Supp. Fig. 3E). Given that PCR typically favors amplification of the short allele (see Fig. 1F), this pattern further substantiates that splicing from *extendedHTT* to *HTT* exon 1 mainly occurs on transcripts that harbor an expanded repeat. Supporting these results, the abundance of *extendedHTT* transcripts in our CAG-enriched short-read sequencing data increased with the repeat length (Pearson’s correlation coefficient, r, = 0.98 between CAG repeat number and *extendedHTT* read count relative to *HTT*, Fig. 3F). Similar correlations were observed using other metrics for *extendedHTT*, like the abundance of splice junctions (Fig. 3F). These repeat-length-dependent RNA processing defects are consistent with prior reports of mis-splicing of *HTT* intron 1 in HD^36^, which we also observed in these cells (Fig. 3A, Supp. Fig. 3F).

To understand why these transcripts have largely escaped detection, we re-analyzed RNA-seq data from 436 LCLs derived from self-reported healthy participants in the 1000 Genomes Project^37^. Splicing of *extendedHTT* to *HTT* exon 1 was exceedingly rare in this population. Only 2 reads (0.02% of the reads spanning the acceptor) reflected these isoforms (Supp. Fig. 3G-H), consistent with our observation that *extendedHTT*-*HTT* exon 1 isoforms are uncommon in the absence of an expanded repeat. Together, these findings establish that *extendedHTT* transcripts splice to the exon 1 of the canonical *HTT* transcript in a repeat-length dependent manner. Although repeat expansion is not strictly required for this splicing, longer repeats increase both the frequency and complexity of these splice isoforms.

### Splicing of *extendedHTT* to the canonical *HTT* exon 1 creates new AUG-initiated open reading frames

We next sought to determine whether *extendedHTT* is translated. Pervasive transcription of the genome generates numerous non-coding RNAs, many of which are nuclear-retained, and thus are not competent for translation^38^. We performed cell fractionation on unaffected and HD LCLs, separating cytoplasmic and nuclear fractions, and quantified the localization of *extendedHTT* by quantitative PCR. *ExtendedHTT* was 39% cytoplasmic (median across 6 cell lines), comparable to canonical *HTT* (median 53% cytoplasmic, Fig. 4A). These numbers are consistent with prior reports showing 20-75% of *HTT* is nuclear retained across various cell lines^39,40^. Reanalysis of ribosome profiling data in GWIPS-viz^41^ from LCLs from apparently healthy individuals (see Methods) revealed elongating ribosome footprints at all common *extendedHTT* exons, albeit at ~1% of canonical *HTT* levels (Fig. 4B). We performed polysome profiling on HD LCLs with various lengths of CAG repeats, and quantified specific RNAs by quantitative PCR (Supp. Fig. 4A, Fig. 4C). *ExtendedHTT* was primarily associated with mono- and polysomes (median 89%, Fig. 4C), similar to mRNAs like *ACTB* (82%) and canonical *HTT* (97%, Fig. 4C). Notably, ribosome association modestly increased with CAG repeat number (Fig. 4C), potentially reflecting a shift toward isoforms with higher ribosome occupancy.

**Figure 4.**
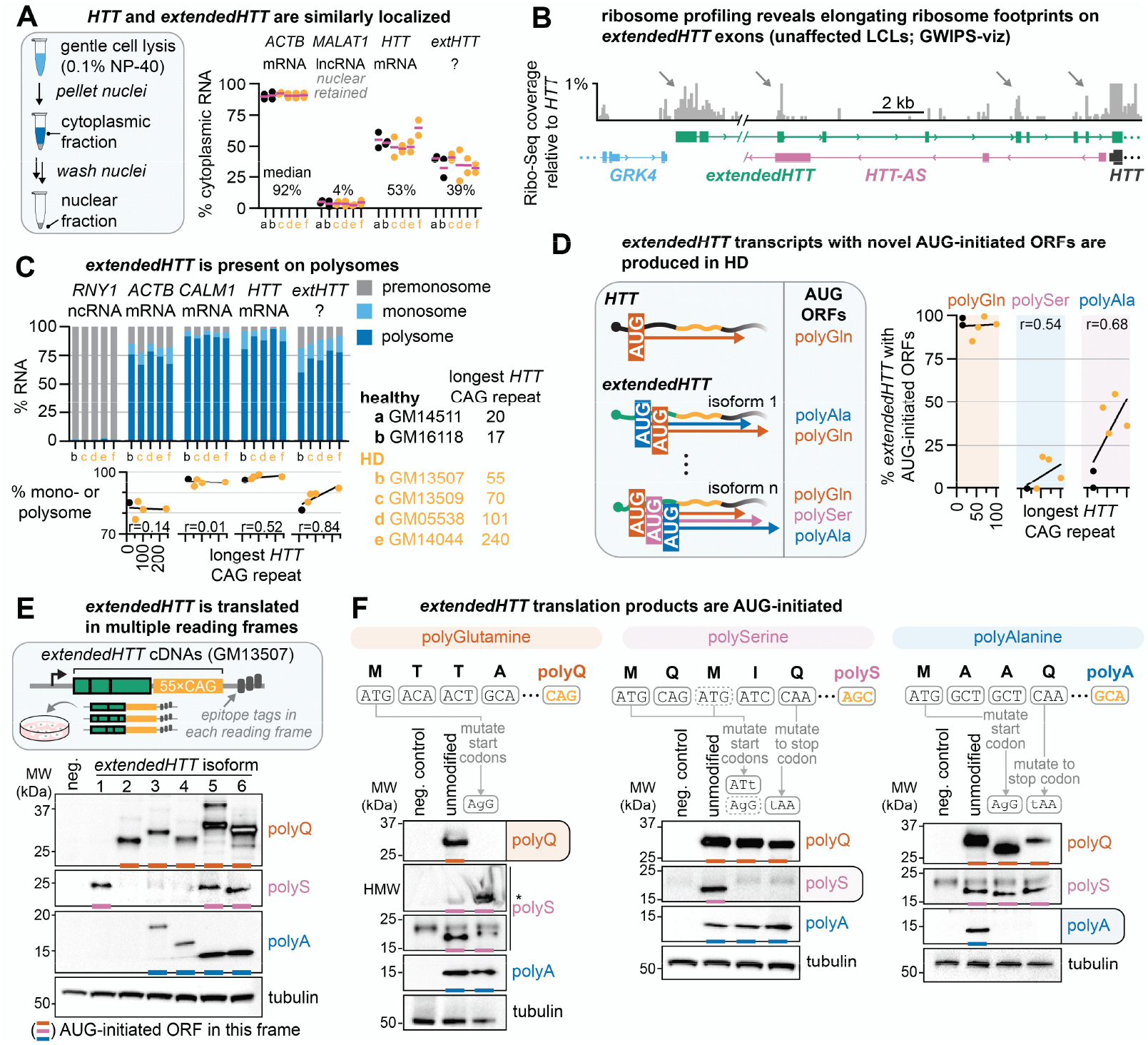
ExtendedHTT transcripts can enable AUG-initiated translation of the CAG repeat in multiple reading frames. **A**. *Left:* schematic for cell fractionation using the REAP protocol^84^. *Right:* Real-time quantitative PCR quantification of the indicated targets, normalized to an *in vitro* transcribed RNA spiked in at equal concentration in each fraction. The housekeeping mRNA *ACTB* and a known nuclear-retained long non-coding RNA (*MALAT1*) are included as controls. Each circle is one biological replicate, with the mean indicated as a purple line. The median of all six cell lines is reported for each target. Sample labels for this panel are shared with panel C. **B**. Re-analyzed ribosome profiling data for elongating ribosomes, obtained from GWIPS-Viz, for unaffected LCLs. Peaks in the sequencing data correlate with *extendedHTT* exons, suggesting that they are engaged with ribosomes. **C**. Real-time quantitative PCR quantification of pooled polysome fractions, normalized to an in vitro transcribed RNA spiked in at equal concentration in each fraction. Highly translated housekeeping RNAs (*ACTB, CALM1*) and a non-coding RNA (human Y-RNA 1; *RNY1*) are included as controls. Data are mean of three replicates. See Supp. Fig. 4A for polysome profiles and pooling. **D**. Quantification of AUG-initiated *extendedHTT* ORFs across the CAG repeat from long-read sequencing (as in Fig. 3B). **E**. *Top:* Schematic for *extendedHTT* ORF reporters transduced into U-2OS cells, depicting the transcription initiation site as a right-facing arrow, with various *extendedHTT* exons fused to *HTT* exon 1, and downstream epitope tags to allow detection of each reading frame. *Bottom:* Immunoblot for the indicated samples. Polyglutamine is directly detected by an anti-polyglutamine antibody, while polyalanine and polyserine are detected by in-frame HA and FLAG epitope tags, respectively. Negative control (neg.) is the parental cell line not expressing an *extendedHTT* transgene. **F**. Targeted point mutations to *extendedHTT* isoform 6 that remove the initiating AUG or insert an immediate stop codon in a given frame abolish the corresponding signal, confirming AUG-initiated translation. *: Loss of the polyglutamine ORF leads to polyserine aggregates that fail to migrate out of the gel well. Immunoblots are representative of ≥ 2 independent experiments.

Next, we asked whether translation of *extendedHTT* could account for spurious out-of-frame proteins. The GENCODE annotated *extendedHTT* isoforms splice directly to exon 2, removing the CAG repeat (Fig. 2B), and thus would not produce repeat-containing proteins. In contrast, the repeat length-dependent *extendedHTT* isoforms we identified retain the repeat tract. We delineated the AUG-initiated ORFs within these isoforms (Supp. Fig. 4B-C, Supplemental Table 2). Surprisingly, every transcript isoform we observed contained AUG-initiated ORFs spanning the CAG repeat, with the majority placing the repeat in the polyglutamine frame (Supp. Fig. 4B-C). These isoforms often introduce N-terminal extensions to the HTT protein (Supp. Fig. 4D). Computational folding analysis suggests that these extensions are not predicted to significantly disrupt HTT folding (Supp. Fig. 4C), although they may affect its localization or function. Splicing also produced numerous transcripts with ORFs in the polyalanine and polyserine frames. Notably, in the polyserine frame, there are multiple stop codons ~30 nt upstream from the repeat (Supp. Fig. 4B). However, when the CAG repeat itself serves as the acceptor, these stop codons are removed, generating AUG-initiated polyserine ORFs (Supp. Fig. 4B-C). In LCLs, up to ~50% of detected isoforms create polyalanine ORFs and up to ~18% create polyserine ORFs, with some transcripts harboring ORFs in all three frames (Fig. 4D, Supp. Fig. 4B-C). Consistent with the observed increase in isoform diversity with increasing repeat length (Fig. 3D), the proportion of transcripts with AUG-initiated ORFs likewise increased with repeat length (Pearson’s correlation coefficient, r, = 0.68 and 0.54 between CAG repeat length and % of transcripts with polyalanine and polyserine ORFs, respectively; Fig. 4D).

The efficiency of translation from an AUG codon is influenced by its surrounding sequence context. The AUG codons in *extendedHTT* occur in relatively modest Kozak contexts (Supp. Fig. 4E). While weak Kozak sequences reduce initiation efficiency, they also enable leaky scanning, allowing ribosomes to initiate at multiple AUGs in different frames^42^. To determine which of the overlapping *extendedHTT* ORFs are translated, we cloned six individual splice isoforms from HD LCL cDNA (GM13507, as in Fig. 3B) containing approximately 55×CAG repeats, each with frame-specific epitope tags downstream of the repeat (Fig. 4D, Supp. Fig. 4F). Upon individually transducing these constructs in U-2OS cells, a variety of translation products were detected (Fig. 4E, Supp. Fig. 4G). Strikingly, the translation products produced from each construct perfectly matched the predicted ORFs for that isoform, even when the transcript harbored multiple overlapping ORFs (Fig. 4E).

To further confirm that these protein products were produced by canonical AUG-dependent initiation, we chose one isoform encoding ORFs in all three reading frames and introduced single-base mutations that either removed the AUG codon(s) or created a stop codon immediately downstream from the relevant AUG. These mutations entirely abrogated translation in that corresponding frame for each of the polyglutamine, polyalanine, and polyserine products (Fig. 4F). Altogether, these findings demonstrate that *extendedHTT* is exported from the nucleus and associates with polysomes in the cytoplasm, where it can undergo AUG-initiated translation to produce repeat-containing proteins in multiple reading frames.

### *ExtendedHTT* is expressed in human brain

Polyalanine and polyserine aggregates have been observed throughout the brain, but particularly in the striatum, the region most heavily affected by neuron death in HD, suggesting they may have a role in disease pathogenesis^7^. Because *extendedHTT* can lead to the production of these proteins, we next sought to determine if *extendedHTT* transcripts are also expressed in brain. We re-analyzed ~182 billion reads from 15 publicly available RNA-seq studies spanning multiple brain regions and disease contexts (excluding HD; see Methods and Supplemental Table 3). Our analysis revealed that *extendedHTT* transcripts were readily detected in this composite dataset at levels comparable to the antisense *HTT* transcript (*HTT-AS*, Fig. 5A, Supp. Fig. 5A). Across all tissues examined, *extendedHTT* transcripts were collectively ~8.2% as abundant as canonical *HTT* by read depth (Supplemental Table 3, Fig. 5A-B). We normalized our data to *HTT* exon 2-6 to avoid biasing our calculations due to the low RNA-seq coverage typically seen across GC-rich exon 1^14^; normalization to exon 1 yielded qualitatively consistent but numerically higher estimates (24% read depth compared to *HTT* exon 1, Supp. Fig. 5B).

**Figure 5.**
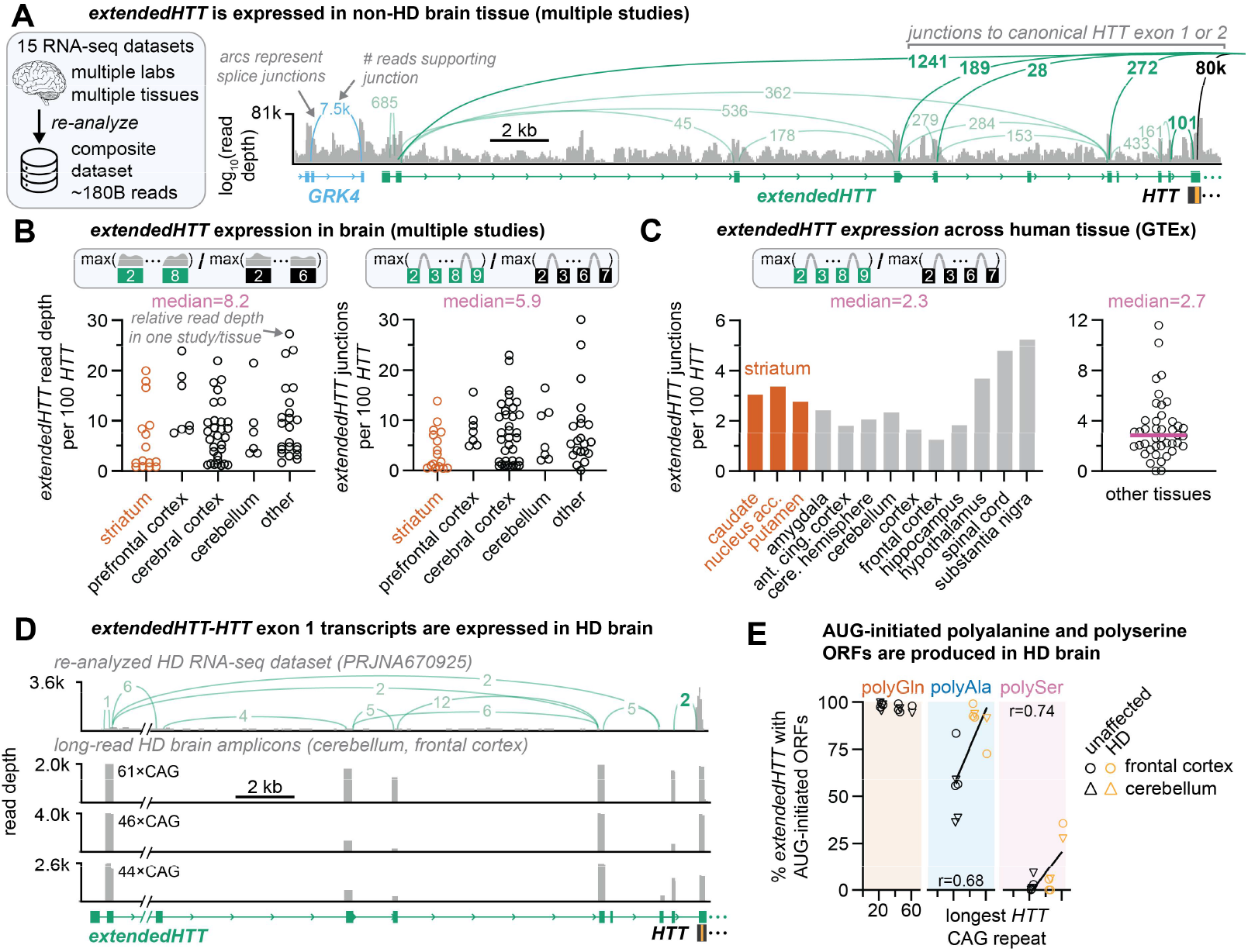
ExtendedHTT is expressed in human brain. **A**. Sashimi plot for composite re-analysis of 15 datasets (multiple studies, tissues but excluding HD; see Methods) reveals reads and splice junctions consistent with *extendedHTT* transcription. Arcs indicating splice junction counts were manually assigned to transcripts by matching splice junctions to transcript annotations. In bold: splice junctions from *extendedHTT* to *HTT* exon 1 or 2, confirming that *extendedHTT* can be joined to the canonical *HTT* exon 1 even in the presumed absence of an expanded repeat. **B**. Quantification of *extendedHTT* abundance in brain by read depth or splice junction count, normalized to *HTT* exons 2-6. Each dot represents a single brain subregion examined in an individual study. Due to variations in study design, these data should be interpreted qualitatively. **C**. Quantification of *extendedHTT* abundance across human tissues in Snaptron’s splicing database for the Genotype-Tissue Expression (GTEx) project. **D**. *Top:* Sashimi plot for reanalyzed RNA-sequencing data from HD patients. Regions corresponding to the *extendedHTT* exons are seen here, providing further evidence that *extendedHTT* is expressed in HD brain. Junctions to *HTT* exon 1 are indicated (*extendedHTT*-*HTT* exon 1 detected in reads SRR12884099.20531474 and SRR12884142.6197957). *Bottom:* Example Nanopore long-read sequencing of cDNA amplicons prepared from CAG-enriched frontal cortex or cerebellum by two-step nested PCR, as in Fig. 3D. **E**. Quantification of AUG-initiated ORFs from long-read sequencing of *extendedHTT* transcripts. Each symbol is one patient/tissue combination from frontal cortex or cerebellum across n=3 unaffected and 3 HD patient brains.

RNA sequencing preparations can contain a small amount of genomic DNA contamination that could potentially contribute to reads across intergenic regions. To definitively confirm that these reads originated from RNA rather than contaminating DNA, we specifically analyzed gapped reads bearing signatures of RNA splicing. Again, *extendedHTT* is expressed across tissues, with splice junctions within *extendedHTT* ~5.9% as frequent as canonical *HTT* junctions in exons 2-6 (Supplemental Table 3, Fig. 5A-B, Supp. Fig. 5B). Notably, we also found >100 reads where *extendedHTT* was spliced to *HTT* exon 1 (0.09% of reads spanning the acceptor; Fig. 5A), at the same splice junctions that we observed in our CAG-enriched HD-LCL datasets. As this composite dataset did not include HD patients, these results suggest that *extendedHTT* transcripts can splice to *HTT* exon 1 in brain, even without documented expansion, albeit infrequently.

While methodological variation across studies precluded quantitative cross-tissue comparisons, *extendedHTT* transcripts were consistently detected in every brain region examined, including the striatum, the primary region of neuronal cell loss in HD (Supplemental Table 3, Fig. 5B). Independent validation using the standardized Genotype-Tissue Expression (GTEx) dataset in the Snaptron splicing database^43^ confirmed that *extendedHTT* transcripts comprise on average ~2.8% of the total *HTT* transcripts detected across all tissues, including all brain regions (median junction abundance 2.3% in brain, Fig. 5C, Supp. Fig. 5C). *ExtendedHTT* levels relative to canonical *HTT* were lower in Snaptron’s larger Sequence Read Archive (SRA) dataset (0.75% junction abundance across 316,000 samples in the Snaptron database; Supp. Fig. 5C-D). Notably, *extendedHTT* expression was 4.7-fold lower in immortalized cell lines compared to other samples (0.17% relative to canonical *HTT* junctions; Supp. Fig. 5D-E). Consistent with this, we could not detect *extendedHTT* transcripts in cell lines such as HEK293T, HeLa, or the embryonic stem cell line isoHD despite relatively deep sequencing (≥ 500 million reads, Supp. Fig. 5F). Although technical factors (e.g., library preparation, sequencing depth, and mapping sensitivity) could contribute, these results suggest that long-term cell culture may suppress *extendedHTT* expression or result in loss of cells expressing these isoforms.

We next examined the prevalence of *extendedHTT* transcripts in existing HD RNA-sequencing datasets. As striatum undergoes extreme atrophy in HD (losing ~95% of the neurons by late-stage disease)^44^ these studies often profile proxy tissues, such as frontal cortex. Despite the inherently poor coverage of the GC-rich *HTT* exon 1 locus, we still detected evidence of *extendedHTT* expression (Fig. 5D). Importantly, we identified spliced *extendedHTT-HTT* exon 1 fusion transcripts using the same splice sites identified in our LCL datasets. In HD frontal cortex sequenced by Labadorf et al., we detected reads supporting these specific splice junctions in two HD patients with modest CAG expansions (~45×CAG, Fig. 5D)^16^.

The low read depth at *HTT* exon 1 in existing datasets limited isoform-level resolution. To directly address this issue, we performed CAG enrichment on RNA from the cerebellum and frontal cortex tissue of HD patients (University of Washington Biorepository, see Methods) and generated RT-PCR amplicons spanning the *extendedHTT*-*HTT* region (as in Figure 3). We identified 46 transcript isoforms in brain that recapitulated the results seen in LCLs (Fig. 5D). Importantly, up to 99% of the detected *extendedHTT* transcripts harbored AUG-initiated polyalanine ORFs, and up to 35% contained polyserine ORFs (Fig. 5E).

Taken together, these analyses establish *extendedHTT* as a bona fide transcript class expressed at ~2-5% of canonical *HTT* across human brain tissues. Our findings demonstrate that, even in the absence of a repeat expansion, *extendedHTT* transcripts can splice into canonical *HTT* exon 1. In HD, increasing repeat length enhances the frequency and diversity of these exon 1 directed isoforms, mirroring our observations in LCLs. These isoforms harbor AUG-initiated polyalanine and polyserine ORFs, thereby providing a mechanistic route to the corresponding repeat proteins.

### Animal models of HD lack the *extendedHTT* promoter region

The impact of aberrant polyserine and polyalanine proteins in disease is yet to be fully defined. Therefore, we asked if animal models could be used to study *extendedHTT*-derived proteins. We first examined the region between *GRK4* and *HTT* that harbors the *extendedHTT* locus in the Multiz alignment of 100 vertebrate genomes to the human genome^45^. This region is conserved between humans and other primates, but is absent in other vertebrates, including those commonly used to model disease, such as mouse (Fig. 6A). Although the distance between human *GRK4* and *HTT* is 34 kb, the equivalent distance in mouse (between *Grk4* and *Htt*, the mouse homologs) is only 6.4 kb. Most of the difference between human and mouse sequences can be explained by the insertion of >20 kb of transposable elements (TEs) between *GRK4* and *HTT* (Fig. 6B). TEs are occasionally exonized and incorporated into transcripts as coding or non-coding exons^46^. *ExtendedHTT* transcripts harbor multiple exonized TEs: exon 5 is derived from an Alu element, exon 6 is a portion of a LINE-1 element, and exons 3, 7, and 8 are fragments of three different ERV1-family retrotransposons (Fig. 6B). Notably, almost all of these TEs are primate-specific (61/66 TEs, Supplemental Table 4).

**Figure 6.**
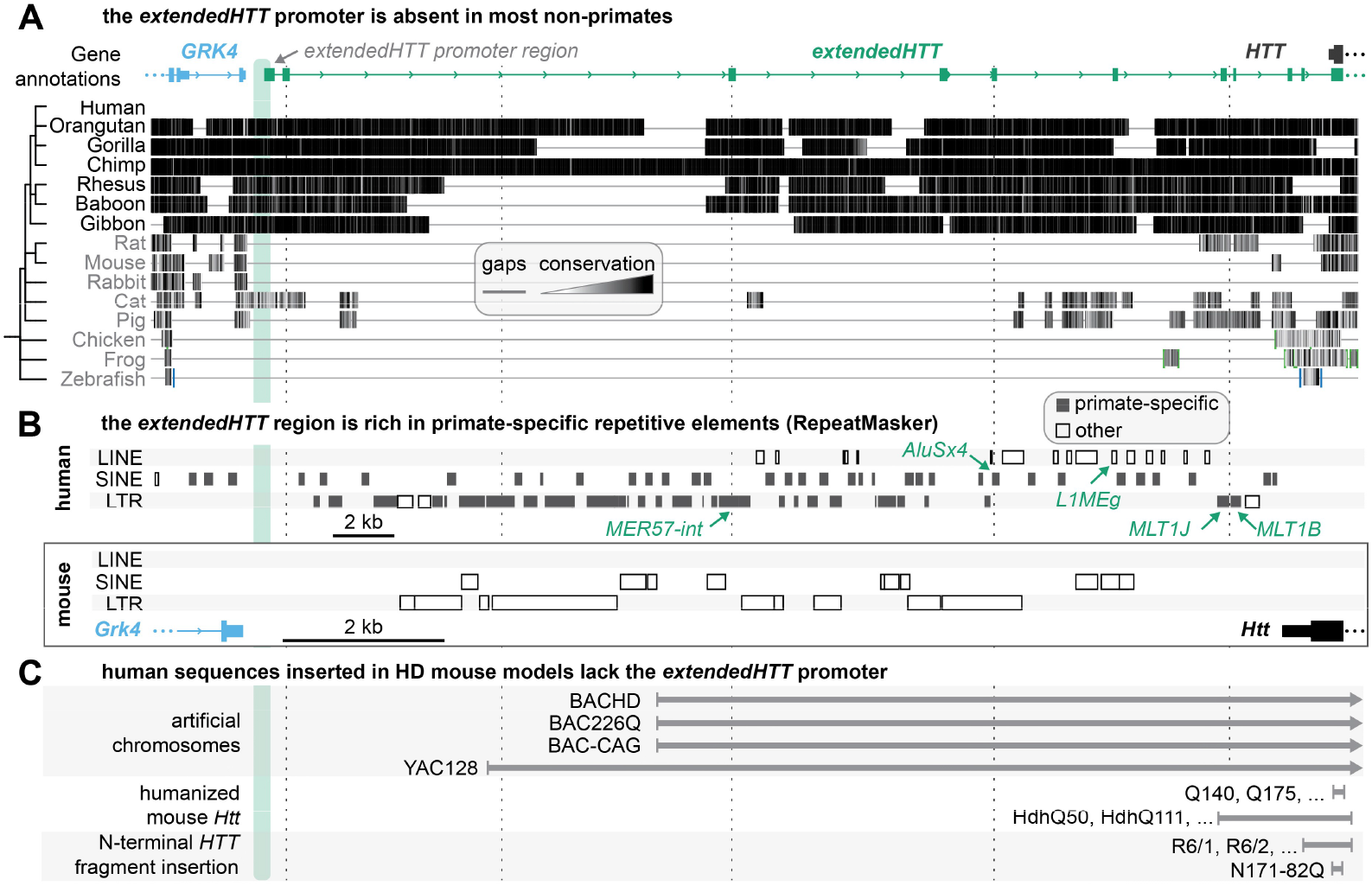
Animal models of HD lack the extendedHTT promoter region. **A**. Pairwise alignment of indicated species to the human genome from Vertebrate Multiz Alignment & Conservation track from the UCSC genome browser. Conservation is indicated by greyscale from white (less conserved) to black (more conserved), and horizontal lines show unaligned gaps. The region likely containing the *extendedHTT* promoter is shaded as a green box, showing that it is absent in mouse, rat, and many other vertebrates. The x-axis is shared with panels B and C, and vertical dashed lines are provided as a visual aid. Phylogenetic tree generated at iphylo.net. **B**. RepeatMasker track from the UCSC genome browser for human and mouse. Primate-specific transposable elements (grey) and elements overlapping with *extendedHTT* exons (green arrows) were manually annotated. **C**. Regions of human sequence included in common mouse models of HD are indicated as grey bars, with rightward arrows indicating additional inserted sequences. Although some models include *extendedHTT* exons, no mouse models include the *extendedHTT* promoter region. See Supplemental Note for further discussion of RAN translation in mouse models.

In addition to missing exons, the mouse *Grk4-Htt* region also lacks homology to the *extendedHTT* promoter region. As a result, humanized mouse models where mouse *Htt* is edited to include an expanded CAG repeat, (such as HdhQ111^47^ or Q140^48^) will lack *extendedHTT* (Fig. 6C). Some mouse models of HD were generated by insertion of bacterial or yeast artificial chromosomes and include fulllength human *HTT* along with portions of the surrounding human chromosome (for example, BACHD^49^, BAC-CAG^9^, and YAC128^50^). However, these models do not extend beyond 25 kb from the 5’ end of *HTT* and thus fall ~8 kb short of the *extendedHTT* promoter region (Fig. 6C). Other commonly used models express short N-terminal fragments of the *HTT* cDNA without substantial genomic context (such as R6/2^51^, Fig. 6C). Altogether, in a survey of HD models (Supplemental Table 5), we did not find any models that include the full *extendedHTT* region along with an expanded repeat. Although several primates have significant homology to the *extendedHTT* region, current primate models of HD were created by semi-random (viral) insertion of small *HTT* cDNAs^52^. Thus, to the best of our knowledge, no animal model of HD includes an expanded CAG repeat in the appropriate genomic context that would allow investigation of the role of *extendedHTT* in disease.

## Discussion

Our work provides a mechanistic explanation for the out-of-frame polyalanine and polyserine proteins in HD. By developing a method to enrich and sequence full-length CAG repeat-containing transcripts, we uncovered a previously uncharacterized, distant promoter for the *HTT* gene. Transcripts originating from this promoter, which we term *extendedHTT*, undergo aberrant, repeat-expansion-dependent splicing to acceptors in the exon 1 region of canonical *HTT* (Fig. 7A). This mis-splicing places the CAG repeat within unexpected, AUG-initiated open reading frames, potentiating canonical translation of the expanded repeat tract in polyalanine and polyserine frames (Fig. 7A). These findings challenge the reliance on non-canonical translation models and demonstrate that expanded repeats can co-opt canonical translation machinery to produce toxic proteins.

**Figure 7.**
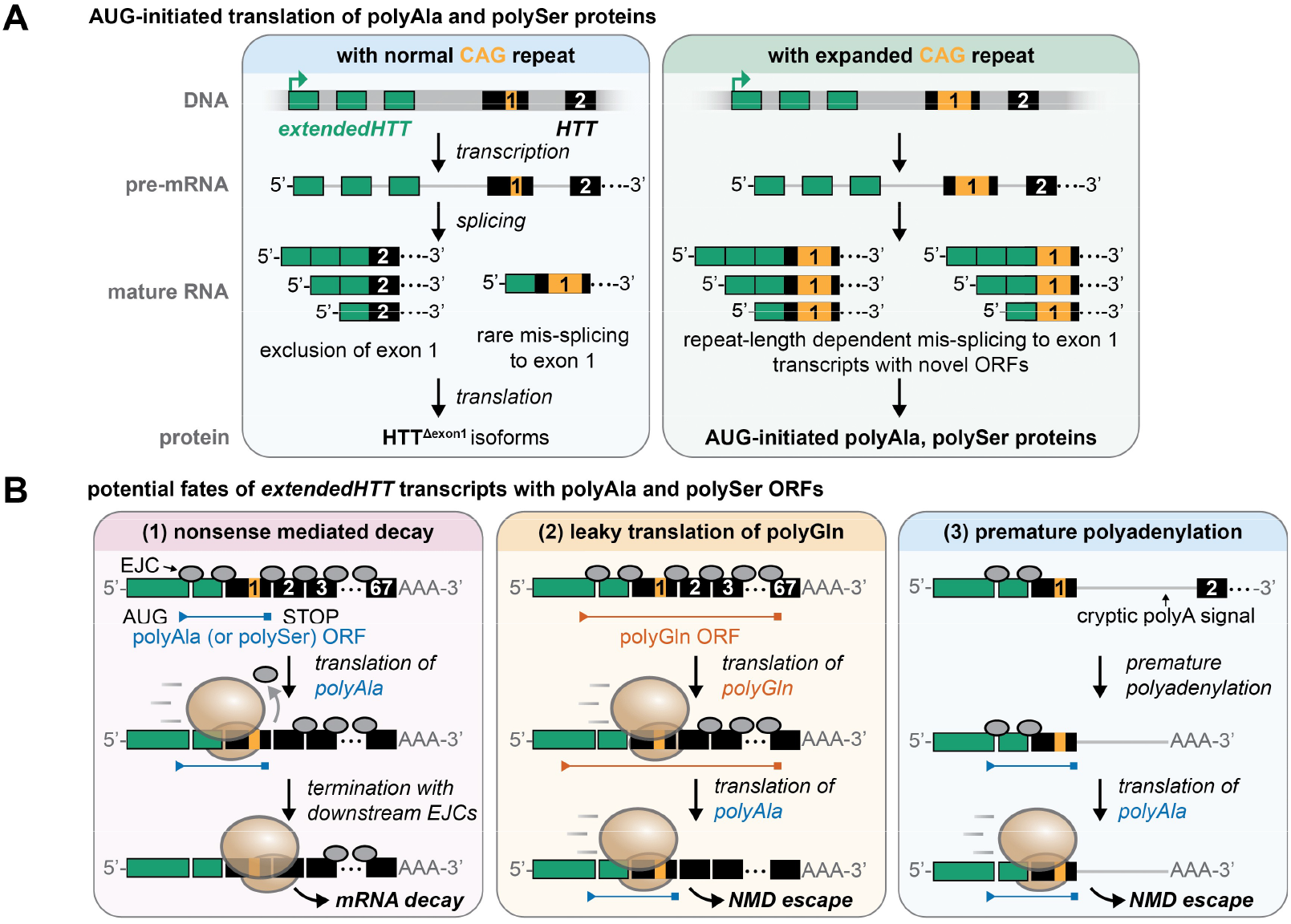
Model for how extendedHTT can contribute to aberrant translation in HD. **A**. In the absence of an expanded repeat, transcripts initiated at the upstream promoter mostly splice to *HTT* exon 2, resulting in the loss of *HTT* exon 1. These transcripts may be translated to produce *HTT*^*Δexon1*^ isoforms, with unknown function. With an expanded repeat, *extendedHTT* transcripts can instead undergo aberrant splicing to *HTT* exon 1, either within the 5’ UTR or to the repeat tract itself, generating multiple *extendedHTT*-*HTT*_*exon1*_ isoforms. All isoforms detected have AUG-initiated open reading frames across the repeat. These isoforms may be canonically translated to produce polyalanine and polyserine, as seen in HD patient brains. **B**. Potential fates of *extendedHTT* transcripts. **(1)** Nonsense-mediated decay (NMD). When translation terminates >50 nt upstream of an exon-exon junction, a residual downstream exon junction complex (EJC) promotes NMD. Thus, mis-spliced *extendedHTT* isoforms with polyalanine and polyserine ORFs that place a stop codon upstream of a junction are therefore expected to be unstable. **(2)** Many of the identified isoforms contain a long, primary ORF in the polyglutamine frame in addition to shorter ORFs for polyalanine and polyserine. Ribosomes translating this main polyglutamine ORF may displace all downstream EJCs, allowing this transcript to bypass NMD. Subsequent scanning/re-initiation can then initiate translation in the polyalanine and polyserine frames. **(3)** NMD escape via premature polyadenylation. Use of a cryptic polyadenylation signal in intron 1 can generate a terminal exon^36,53^, eliminating downstream EJCs relative to the stop codon and thereby evading NMD.

The polyalanine and polyserine ORFs in *extendedHTT* contain termination codons upstream of an exon junction: a canonical trigger for rapid degradation by nonsense-mediated decay (NMD; Fig. 7B). However, their detection in our steady-state RNA-seq implies they must somehow evade cellular quality control. Interestingly, the most abundant isoforms detected in our analysis contain a long ORF in the canonical polyglutamine frame in addition to shorter polyalanine/polyserine ORFs. During early rounds of translation, a ribosome translating this primary polyglutamine ORF could displace the downstream exon junction complexes, effectively allowing the transcript to bypass NMD (Fig. 7B). Once licensed, the same transcript can potentially undergo leaky scanning or re-initiation at alternative AUGs to produce polyalanine and polyserine proteins. In addition, *extendedHTT* transcripts can also be prematurely polyadenylated at cryptic intronic sites (such as in intron 1)^36,53^, which eliminates downstream EJCs and likewise allows escape from NMD (Fig. 7B).

*Why has extendedHTT escaped detection so far?* Three interconnected factors explain why *extendedHTT* remained undetected. First, is the technical challenge: from our analysis, *extendedHTT* may be expressed at levels 2-5% of canonical *HTT* (Fig. 5). Compounding this challenge, the GC content and secondary structure of both canonical *HTT* exon 1 and upstream *extendedHTT* exons impair reverse transcription and sequencing coverage of these regions (Supp. Fig. 5G). We also observed that the libraries prepared with methods that deplete rRNA (rather than enriching polyA+ mRNA) showed reduced levels of *extendedHTT* (Supp. Fig. 5E). Second, the *extendedHTT* promoter appears to be inactive in many commonly used cell lines, including neuronally-differentiated stem cells (Supp. Fig. 5F). These cell lines also do not express the annotated antisense *HTT* transcript (*HTT-AS*) that is detected in brain (Supp. Fig. 5A). This lack of expression of both *extendedHTT* and *HTT-AS* may indicate tissue-specific expression that is not captured by cell lines, or imply that cells expressing these transcripts were selected against during cell-line establishment. Third, the *extendedHTT* locus is primate-specific, likely arising from insertions of transposable elements. Consequently, no existing animal models of HD, including humanized BAC or YAC transgenic mice, contain the necessary genomic architecture to produce these transcripts, creating a critical blind spot in our study of these transcripts and their contribution to disease (see Supplemental Note for additional context on RAN translation in mouse models).

Although *extendedHTT* is less abundant than canonical *HTT*, low transcript frequency does not necessarily limit pathogenic potential. The *HTT*_exon1a_ transcript, generated through alternative polyadenylation, is similarly rare and difficult to detect by RNA sequencing, yet it yields toxic N-terminal *HTT* fragments readily observed in HD brain that are central to disease pathology^36,53^. Similarly, while we detect relatively few *extendedHTT* transcripts, the polyserine and polyalanine products they encode have been documented in HD brains^7^. Notably, recent work suggests that polyserine peptides can be particularly pathogenic: they can nucleate tau aggregation and promote cross-seeding between pathological proteins^54^. In long-lived, post-mitotic cells like neurons, even a low level of continuous production of aggregation-prone proteins can lead to significant accumulation and toxicity over time. Moreover, slow protein turnover in the brain^55^ and incomplete proteasomal^56^ or autophagic clearance^57,58^ of repetitive sequences could further amplify pathogenic potential.

Several factors could increase the frequency or impact of *extendedHTT in vivo*. Genetic variation can tune splice decisions; for example, the SNP rs146151652 increases the strength of a principal splice acceptor within exon 1, potentially modulating splicing from *extendedHTT*. Somatic instability changes the CAG repeat number across tissues and with age^3^, creating cellular mosaicism in which the *HTT* tract can expand to hundreds, and in rare cells, even thousands, of repeats^20,36,59–64^. We find that longer tracts are more prone to aberrant processing, which may raise both the frequency and the diversity of repeat-containing *extendedHTT* splice isoforms in the neuronal populations most vulnerable in HD. Furthermore, several *extendedHTT* isoforms place the repeat in the polyglutamine frame but fuse it to novel N-terminal sequences that are distinct from the canonical HTT protein. The amino acid sequences flanking a polyglutamine tract are critical determinants of its folding and aggregation propensity^65^ and these new sequence contexts may fundamentally alter the properties of the resulting HTT protein. While the full toxic potential of these aberrant proteins remains unresolved, the disease-associated splice junctions provide a precise therapeutic handle: splice-switching ASOs^66^ targeting the *extendedHTT/exon 1* junctions could potentially block aberrant protein production while sparing canonical *HTT*.

Besides HD, out-of-frame translation products have been reported at multiple repeat-expansion loci and are often discussed under the broad umbrella of “RAN translation”^8^. Several mechanisms have been identified to contribute to these products including translation initiation at near-canonical start sites (ACG in fragile X tremor ataxia syndrome^67^ and CUG in *c9orf72*-mediated amyotrophic lateral sclerosis^68^) and ribosomal frameshifting within expanded repeats^68–72^. Additionally, the secondary structures arising from expanded GC-rich repeats may act as internal ribosome entry sites to allow cap-independent (and AUG-independent) translation within the repeat itself^4,73–75^. In previous work, we showed that aberrant splicing of the repeat-containing transcript followed by canonical translation initiation can account for some instances of RAN translation^12^. Here, we build on this model, and provide evidence that aberrant splicing can generate transcript variants in HD that reposition the CAG repeats into alternative AUG-initiated frames, providing a canonical pathway for the production of seemingly out-of-frame proteins.

This splicing-mediated generation of alternative ORFs may extend to other repeat-expansion disorders in which “RAN” products are reported. In several cases, RNAs with pathogenic repeat expansions exhibit prolonged nuclear retention^76^, thus increasing opportunities for the spliceosome to engage weak or cryptic splice sites^77^. Consistent with this, intronic expansions of CCUG and GGGGCC in myotonic dystrophy type 2 and amyotrophic lateral sclerosis have been shown to promote intron retention, converting the intronic repeat into a new exon^78,79^. Likewise, splicing may potentially reposition repeats within AUG-initiated reading frames that provide a canonical route to out-of-frame proteins. We note that this model is not mutually exclusive with other proposed mechanisms, and different routes may dominate depending on disease, cell type, and state.

More broadly, many pathogenic repeat loci are situated in intricate genomic neighborhoods with alternative promoters, antisense transcripts, and extensive isoform diversity^80–82^. This complexity is further enhanced upon repeat expansions, which can remodel local chromatin architecture and influence TAD boundaries^83^. These regions are typically GC-rich, repetitive, and highly structured, causing them to be systematically under-represented in standard RNA-seq datasets. As a result, the full repertoire of transcripts produced in disease often remains incompletely defined. The affinity purification technique implemented here is readily generalizable to other repeat expansions and provides a powerful tool to directly address this issue. By enabling the targeted enrichment of full-length, repeat-containing transcripts, this approach allows for an unbiased characterization of the complete set of transcripts arising from these complex loci, revealing both disease mechanisms and potential therapeutic targets.

## Supporting information

Supplemental Table 7 - Oligonucleotides

Supplemental Table 1 - extendedHTT exons

Supplemental Table 2 - extendedHTT transcripts

Supplemental Table 3 - extendedHTT in brain

Supplemental Table 4 - extendedHTT TEs

Supplemental Table 5 - HD animal models

Supplemental Table 6 - Constructs

## Acknowledgements

We thank the members of the Jain lab for helpful discussions. This work was supported by grants from the NIH (R00AG053434), Chan Zuckerberg Initiative, David and Lucile Packard Foundation, and the Smith Family Awards Program. This paper was typeset with the bioRxiv word template by @Chrelli: www.github.com/chrelli/bioRxiv-word-template

## Author contributions

R.A. and A.J conceived this study, designed experiments, and interpreted results. R.A. performed most experiments and data analysis. T.S. assisted with optimizing repeat-affinity RNA purification experiments. R.A. and A.J. wrote the paper with contributions from all authors.

## Methods

### Cell culture

HEK293T (CRL-3216; RRID:CVCL_0063) and U-2OS (HTB-96; RRID:CVCL_0042) were obtained from ATCC, lymphoblastoid cell lines (unaffected: GM14511, GM16118; HD: GM06685, GM04856, GM13507, GM13509, GM05538, GM14044; other CAG repeat expansions: GM23709, GM03562, GM14982, GM06926) were obtained from the Coriell Institute. Cell lines were tested for mycoplasma at least quarterly using a PCR-based assay^85^. Lymphoblastoid cell lines were maintained at 37 °C in 5% CO_2_ in Roswell Park Memorial Institute 1640 Medium (RPMI; Gibco 11875135) supplemented with 15% cosmic calf serum (CCS; HyClone SH30087.04) and 1% Penicillin-Streptomycin-Glutamine (PSG; Gibco 10378016). HEK293T and U-2OS cells were maintained in Dulbecco’s Modified Eagle Medium (DMEM; Gibco 11965126) with 10% CCS and 1% PSG. Adherent cells were passaged 3 times per week at 1:10 dilution. LCLs were passaged by 1:4 dilution as needed to maintain 200-800×10^3^ cells/mL, typically every 2-3 days.

### Organisms/Strains

Plasmids were propagated in Stbl3 E. coli (Invitrogen C737303) grown at 37°C in Luria-Bertani (LB) medium. All plasmids were verified by Oxford Nanopore sequencing (Quintara Biosciences).

### Cloning and plasmid generation

Complete sequences for all plasmids used in this study are provided in Supplemental Table 6. CAG reporter plasmids with varying number of repeats were previously described^13^. pCMV-VSV-G was a gift from Bob Weinberg (Addgene plasmid #8454)^86^. psPAX2 was a gift from Didier Trono (Addgene plasmid #12260). *ExtendedHTT* cDNA reporters were first amplified from CAG-enriched GM13507 cDNA using nested PCR (described below), after which adapters were added for Gibson assembly into an MluI and NotI digest of a CAG reporter plasmid. Mutations to the *extendedHTT* ORFs were introduced by PCR to prepare two fragments that were inserted between MluI and NotI sites by Gibson assembly. See Supplemental Table 7 for oligonucleotide sequences.

### Repeat-affinity RNA purification and short-read sequencing

For large-scale RNA affinity purification, LCLs were expanded to ~100×10^6^ cells, then pelleted at 500×g and snap-frozen. Cell pellets were stored at - 80 °C until RNA isolation by TRIzol reagent (Invitrogen 15596018) according to the manufacturer’s protocols. We typically recovered ~10 μg RNA per 10^6^ LCL cells. In a typical large-scale RNA affinity purification, 500 μL of RNA (300-1500 μg total RNA) was mixed with 5 μL 1 M DTT, 10 μL SuperaseIn (AM2694), 1 μL 100 μM biotinylated bait oligonucleotide, and 516 μL 2× hybridization buffer (50% v/v formamide [Invitrogen AM9342], 2 mM EDTA [Invitrogen AM9010], 0.2% sodium dodecyl sulfate, 9.6X saline-sodium citrate buffer [SSC; Invitrogen 15557044]). RNA was hybridized to the bait oligos at 37 °C for 90 minutes on a rotary mixer, then 300 μL streptavidin beads (NEB S1421S) pre-equilibrated in 1x hybridization buffer were added and incubated at 37 °C for 30 minutes to capture bound RNAs. For gentle wash conditions, the beads were washed with 1 mL of 1× hybridization buffer four times at room temperature (RT). For stringent wash conditions, the beads were washed twice with 1× hybridization buffer at RT, then twice washed at 60 °C for 30 seconds in stringent wash buffer (1× SSC, 25% formamide), with the wash buffer removed while holding at 60 °C in a thermocycler. To degrade residual genomic DNA, we treated the beads with Turbo DNase (Invitrogen AM2238; 50 μL reaction, containing 5 μL 10X DNase buffer, 1 μL Turbo DNase, 1 μL SuperaseIn, and 1 μL 100 mM DTT) at 37 °C for 20 minutes. The DNase buffer (containing a small amount of eluted RNA) was saved, and the remaining RNA was eluted from the beads with 50 μL 95% formamide buffer (95% formamide, 10 mM EDTA, 10 mM Tris pH 8) at 60 °C for 5 minutes. The DNase buffer and eluted RNA were pooled and cleaned up with Zymo RNA Clean & Concentrate columns (R1016) per manufacturer instructions. Following internal quality control by qPCR (see Supp. Fig. 1D), polyA-selected libraries were prepared by Novogene Corporation and sequenced on a NovaSeq 6000 (Illumina) as 150bp paired-end reads to a depth of at least 25M reads per sample.

### Short-read RNA-sequencing analysis

Low quality read ends and sequencing adapters were removed from the reads using cutadapt^87^ (version 3.7) with commands “-a AGATCGGAAGAG --error-rate=0.1--times=1 --overlap=5 --minimum-length=20 --quality-cutoff=20”. The processed reads were aligned to hg38 using STAR^88^ (2.7.1a) with default arguments. Duplicate reads were removed using samtools^89^ markdup (1.11). Short-read transcript assembly was performed with StringTie^35^ (version 2.2.1) with default options. Coverage across gene bodies was calculated using RSeQC^90^ (5.0.1). Per-gene coverage was quantified by featureCounts^91^ (version 1.6.2).

### Analysis of public datasets

#### Transcript annotations

GENCODE annotations for the *extendedHTT* transcripts were provided by the HAVANA project. As of GENCODE v43, HAVANA’s manual annotation remark files lacked entries for *extendedHTT* transcripts, implying that they were included by an automated system. One automated system in use is APPRIS^92^, which integrates RNA and protein evidence to evaluate transcripts. APPRIS scores one *extendedHTT* transcript (ENST00000680956) as alternative:2, reflecting an alternative (non-primary isoform) transcript model that is conserved in fewer than three non-primate species, consistent with our findings that *extendedHTT* is absent in non-primates (see Figure 6).

#### Re-analysis of public datasets

annotated transcription start sites (TSS) from refTSS; 5’ TSS tags (CAGE, Fantom5, RAMPAGE); promoters from the EPDnew database; repetitive elements from RepeatMasker; and Multiz 100-way vertebrate annotations were exported from their respective UCSC Genome Browser tracks for hg38. ChIP-Seq and chromatin accessibility were exported from ChIP-Atlas including all cell types. Ribosome profiling data were aggregated from all LCL experiments in the GWIPs-viz database for elongating ribosomes.

#### Brain RNA-seq re-analysis

We non-exhaustively searched the NCBI Sequence Read Archive for RNA-sequencing datasets that were prepared with traditional Illumina-type library preparation kits; that examined at least one region of human brain across multiple subjects; and were sequenced with 75 bp or longer paired-end reads. We used a custom pipeline that aligned the dataset to hg38 using STAR. We collected all reads mapping within 200 kb of the *HTT* gene using samtools, and combined all reads from a dataset (comprised of multiple samples) into a single bam file. In total, we analyzed 15 datasets totaling 182 billion reads (PRJEB14594, PRJNA393104, PRJNA434002, PRJNA512012, PRJNA527986, PRJNA563037, PRJNA600414, PRJNA644618, PRJNA673437, PRJNA753141, PRJNA753516, PRJNA756274, PRJNA780522, PRJNA836496, PRJNA845531). Coverage per-exon was determined using samtools coverage, and splice junctions were counted with samtools. See extract-brain-coverage.sh and brain-re-analysis.ipynb for coverage and junction counting. Relative levels of *extendedHTT* (by coverage or splicing) were calculated as:

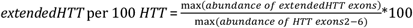

We used the maximum value for these calculations due to the high heterogeneity in *extendedHTT* exon inclusion, and potential reduced coverage of exons due to GC content and secondary structure. See Supplemental Table 3 for per-sample counts and our calculations for splice junctions and read depth. Sashimi plots were generated by IGV and hand-curated to assign splice junctions to genes, highlighting *extendedHTT* and *HTT* splice junctions supported by ≥25 reads.

#### Snaptron

We queried the Sequence Read Archive (SRA; srav3h), Genotype-Tissue Expression (GTEX; gtexv2), and The Cancer Genome Atlas (TCGA; tcgav2) compilations in the Snaptron splicing database^43^ for splice junctions in the *GRK4*-*HTT* region, e.g., “https://snaptron.cs.jhu.edu/srav3h/snaptron?regions=chr4:3037390-3250000”.

Splice junctions matching our identified *extendedHTT* exons and canonical *HTT* exons were included in the analysis, using the same methods described above. See snaptron_queries.ipynb for code.

### Long-read RNA-sequencing analysis

For LCL amplicons, reads were aligned to the *GRK4*-*HTT* region with the CAG repeat edited to 200×CAG, using minimap2^93^ (version 2.24-r1122) and options “-x splice -u b --secondary=no” For *extendedHTT* cDNA reporters, reads were instead aligned to the appropriate plasmid map.

#### Repeat length analysis

We first selected reads that aligned to regions on both side of the *HTT* CAG repeat, excluding those that truncated within the repeat. Sizing was performed using custom python code (see short-read-analysis.ipynb and long-read-analysis.ipynb) that attempted to locate the largest CAG tract in the read, allowing up to 3 errors (insertions, deletions, or substitutions). This script was validated using several LCL cell lines with known *HTT* repeat lengths, then used to infer the repeats present in other samples.

#### Isoform analysis

After alignment, we selected full-length reads that span from primer-to-primer. Because exponential amplification of expanded repeats can introduce truncations in the repeat, reads had a distribution of repeat lengths that complicated isoform analysis, precluding detection of the CAG repeat tract as the splice acceptor. We overcame this in two ways, which yielded equivalent results (see Supp. Fig. 3D). First, we used SATCfinder to trim repeats from reads, as previously described^12^. Second, we post-processed the reads to standardize the reads to carry identical read lengths using a custom python script (long-read-analysis.ipynb). Transcript assembly was performed using StringTie with options “--rf -f 0.005 -L -E 25”. These settings include any isoform making up at least 0.5% of isoforms; because all samples had at least 2000 reads, only isoforms supported by ≥10 reads were included in our analysis.

### Real-time quantitative and endpoint PCR

RNA was isolated from approximately 10^7^ lymphoblastoid cells or 10^6^ U-2OS cells using TRIzol according to the manufacturer’s protocols. Contaminating genomic DNA was removed using ezDNase (Invitrogen 11766050), followed by reverse transcription using SuperScript IV VILO master mix (Invitrogen 11756050). Briefly, 1 μg total RNA in a 5 μL volume was incubated with ezDNase at 37 °C for 2 minutes. This reaction was diluted with 3 μL nuclease-free water before addition of 2 μL SuperScript IV VILO master mix. The mixture was incubated at 25 °C for 10 minutes to anneal primers, 50 °C for 20 minutes for reverse transcription, followed by inactivation at 85 °C for 5 minutes.

For quantitative PCR, a 1:100 dilution of the reverse transcription reaction was quantified with SYBR Green PCR Master Mix (Applied Biosystems 4309155) and 200 nM primers using the QuantStudio 3 RT-PCR system (Applied Biosystems A28567).

To generate LCL amplicons for long-read sequencing, we found nested PCR was required to avoid off-target amplification, as the 3’ end of *HTT* exon 1 lacks unique primer binding sites. In nested PCR, cDNA prepared from CAG-enriched RNA (corresponding to approximately 0.5 μg of total RNA) was amplified in a 25 μL reaction with Advantage GC 2 polymerase (Takara Bio 639114) using 1 M GC Melt and 200 nM primers. The reactions were cycled 20 times with outer PCR primers, with denaturation at 94 °C for 10 seconds, annealing at 58 °C for 10 seconds, and extension at 68 °C for 90 seconds. At completion, 1 μL of thermolabile exonuclease I (NEB M0568L) was added and primers were degraded at 37 °C for 20 minutes, followed by heat inactivation at 65 °C for 1 minute. Inner PCR primers were added and the reaction was topped up to 50 μL, after which a further 20 cycles of PCR performed. Amplicons for the *extendedHTT* cDNA reporters were prepared similarly, but with only one round of 25-cycle PCR. The amplicons were cleaned up with Zymo DNA Clean & Concentrate kit (D4004) and Oxford Nanopore sequencing performed by Plasmidsaurus. See Supplemental Table 7 for oligonucleotide sequences.

### Western blot

Cells were washed with DPBS and lysed with 160 μL RIPA buffer (25 mM Tris-HCl pH 7.5 (Invitrogen 15567027), 150 mM NaCl (Invitrogen AM9760G), 1% (v/v) NP-40 (Fisher Scientific AAJ19628AP), 1% (w/v) sodium deoxycholate (Sigma-Aldrich D6750), 0.1% (w/v) SDS (Bio-Rad 1610302)) supplemented with 1% (v/v) HALT protease and phosphatase inhibitors (Thermo Scientific 78429) and 125 U/mL Benzonase nuclease (EMD Millipore E1014). The cell lysate was homogenized by passing through a 21-gauge needle (Becton Dickinson 305110) 5 times, and incubated on a nutator for 30 minutes at 4 °C. We noticed that aggregation-prone poly-glutamine, -serine, and -alanine proteins were frequently lost in the pellet fraction after centrifugation at 21000×g or even at 3000×g for 5 minutes. Thus, homogenized lysate was used without further clean-up for electrophoresis. Lysates were incubated with NuPAGE 4× LDS buffer (Invitrogen NP0008) with 50 mM dithiothreitol (DTT; Thermo Scientific R0861) at 70 °C for 5 minutes for most assays. When probing polyalaninecontaining proteins, lysates were instead heated at 37 °C for 10 minutes, which we previously found to reduce aggregation (see Supp. Fig. 5O in ^12^). Samples were separated on a Bolt 4-12% Bis-tris polyacrylamide gel (Invitrogen NW04122) run at 200 V for 28 minutes, then transferred to PVDF membranes (Invitrogen IB24002) using the iBlot 2 dry blotting system (Invitrogen IB21001). Membranes were blocked in 5% (w/v) skim milk (BD Biosciences 232100) in TBST (tris-buffered saline (Fisher Scientific AAJ60764K3) with 0.1% (v/v) Tween-20 (Fisher Scientific BP337) for one hour at room temperature, then incubated with primary antibodies in 1% (w/v) skim milk in TBST at 4 °C overnight. The membranes were washed 4 times for five minutes each with TBST, then incubated with the appropriate secondary antibody in 1% (w/v) skim milk for one hour at room temperature. After 4 five-minute washes in TBST, chemiluminescence was detected using SuperSignal West Femto Maximum Sensitivity Substrate (Thermo Scientific 34095) with a ChemiDoc XRS+ imager (Bio-Rad). Primary antibodies were used at the following dilutions: mouse anti-FLAG (1:1000, Sigma-Aldrich F1804), mouse anti-HA (1:1000, BioLegend 901501), mouse anti-polyQ (1:20,000, Sigma-Aldrich MAB1574), and rabbit anti-β-tubulin (1:2000, Cell Signaling Tech. 2146). Secondary antibodies: goat anti-rabbit HRP conjugate (1:2000, Sigma Aldrich A0545), rabbit anti-mouse HRP conjugate (1:2000, Sigma Aldrich A9044).

### Cell toxicity assays

∼10,000 cells were plated per well in a 6-well plate. The next day, the media was replaced with DMEM with FBS and PSG supplemented with or without doxycycline. After five days, when the cells were approximately 75% confluent, the supernatant containing floating (dead) cells was collected. Cells were washed with DPBS to collect loosely attached cells, and the wash added to the supernatant. The adherent cells were trypsinized and added to the supernatant. Cells were pelleted at 500×g for 3 minutes, then resuspended in 200 μL DMEM. Dead cells were stained with trypan blue (Invitrogen T10282). The number of live cells was quantified for ≥ 3 technical replicates using a Countess II FL Automated Cell Counter (ThermoFisher AMQAf1000). The cell count for doxycycline induction was normalized to the without-doxycycline condition for each of three biological replicates per condition.

### Polysome profiling

20 million LCL cells were washed once with DPBS and then lysed in 1 mL lysis buffer (20 mM HEPES pH 7.5, 100 mM KCl, 5 mM MgCl2, 0.1% Triton-X-100, 2 mM DTT, 100 μg/mL cycloheximide [(Sigma Aldrich C1988-1G], 20 μ/mL RNasin Plus [Promega N2615], supplemented with cOmplete protease inhibitor [Roche 11836170001]). Lysates were incubated on ice 10 minutes, homogenized 5× with a 27g needle, and centrifuged at 1500×g for 10 minutes at 4 °C. 300 μL of each supernatant was loaded onto a 10-50% sucrose gradient (supplemented with 20 mM HEPES pH 7.5, 5 mM MgCl2, 100 mM KCl, 10-50% (w/v) sucrose, 100 μg/mL cycloheximide, 20 u/mL RNasin Plus, and 2 mM DTT), centrifuged in a Beckman ultracentrifuge (rotor SW41Ti) at 36000 rpm for 2 hours at 4 °C, and then fractioned on a BioComp gradient fractionator.

We mixed 100 μL of each fraction in the pre-monosome (4 fractions), monosome (3 fractions), and polysome (6 fractions) regions, padding up to 600 μL with 10% sucrose buffer as needed. To allow for normalization between fractions, 5 ng of an *in-vitro* transcribed nanoLuc RNA (MEGAscript T7 Transcription Kit, Invitrogen AMB13345, prepared as manufacturer’s protocol) was added to each pooled fraction. After pooling fractions, 600 μL TRIzol (Invitrogen 15596018) and 240 μL chloroform was added, and the fractions mixed aggressively for 15 seconds, then incubated at room temperature for 3 minutes. After centrifuging at 12000×g for 5 minutes at 4 °C, the upper phase was added to 1.2 mL of isopropanol with 2 μL GlycoBlue (Invitrogen AM9516), mixed well, and incubated overnight at -20 °C. After centrifuging at 12000×g for 10 minutes at 4 °C, the supernatant was discarded and the RNA pellet washed with 1 mL of 75% ethanol. Following centrifugation at 7500×g for 5 minutes at 4 °C, the supernatant was discarded and the pellets allowed to air dry for 5 minutes before re-suspending in 16 μL of 60 °C RNase-free water. 4 μL of SuperScript IV VILO Master Mix was added to the entire RNA isolate, which was reverse transcribed as described above.

## Supporting Information

### Supplemental Note

In this work, we show that the *HTT* gene harbors an upstream promoter (Fig. 2), giving rise to *extendedHTT* transcripts that undergo repeat-length dependent missplicing (Fig. 3). Moreover, *extendedHTT* transcripts can generate AUG-initiated polyserine and polyalanine proteins (Fig. 4). However, *extendedHTT* is absent in mouse models of HD, which lack the promoter region (Fig. 6). This raises the question: *why do some models nevertheless display translation of polyalanine and polyserine proteins?* Here, we address this question by examining the two models (BAC-CAG and N171-82Q) in which these proteins have been detected. In both cases, intronic sequences upstream of the expanded CAG repeat are present, suggesting that RNA mis-processing may again facilitate the production of polyalanine and polyserine proteins.

The BAC-CAG model was generated by insertion of a bacterial artificial chromosome (BAC; RP11-866L6) containing full-length *HTT* and surrounding genomic context^1,2^ (Supp. Note A). To determine the sequences retained after integration, we re-analyzed RNA-seq data^2^ from BAC-CAG and WT controls, aligning reads to the mouse genome (mm39) as well as to human *HTT* with 150-kb of flanking sequence. As expected, BAC-CAG has RNA-seq coverage across *HTT*, but also over regions ~7.5 kb upstream and ~60 kb downstream of the gene (Supp. Note B). These regions exhibit splicing, indicating that they are expressed as RNAs, and are absent from WT mice, consistent with their derivation from the BAC insertion (Supp. Note B). The upstream region contains *extendedHTT* exons 7-10 (lacking a promoter), while the downstream region encompasses *MSANTD1* and the promoter and first exon of *RGS12* (Supp. Note B).

For each RNA target, the percent of RNA in each pooled fraction was calculated by a previously described method^94^:

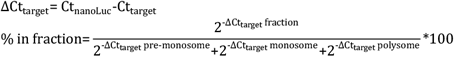

### Cell fractionation

REAP was performed according to previous literature^84^, with minor modifications for isolating RNA. In brief, 2 million cells were pelleted and resuspended in 400 μL of ice-cold PBS, then transferred to a microcentrifuge tube and centrifuged at 15000×g for 10 seconds. The supernatant was discarded and 200 μL of cytoplasmic extraction buffer (0.1% NP-40 in PBS) was added. The whole-cell pellet was gently resuspended by pipetting 5x with a P1000. After centrifugation at 15000×g for 10 seconds, the supernatant (cytoplasmic fraction) was added to 1 mL of TRIzol (Invitrogen 15596018) containing 5 ng of an *in-vitro* transcribed nanoLuc RNA to allow for normalization between fractions. The nuclei pellet was washed once with 1 mL of ice-cold cytoplasmic extraction buffer and pipetted 10× with a P1000, then centrifuged at 15000×g for 10 seconds. After discarding the supernatant, 200 μL of cytoplasmic extraction buffer was added and the fraction containing nuclei (and other intact organelles, like mitochondria) was transferred to 1 mL of TRIzol with 5 ng of nanoLuc RNA.

RNA was isolated from TRIzol according to manufacturer’s instructions as described for polysome profiling. Following resuspension in 20 μL of RNase-free water, typical yields were around 500 ng/μL for cytoplasmic fractions, and 200 ng/μL for nuclear fractions. 2 μL of this RNA was treated with ezDNase and used as template for reverse transcription with SuperScript IV VILO master mix and analyzed by quantitative PCR, as described above.

The BAC-CAG mouse has been annotated to carry two tandem BAC insertions^2^. This arrangement could position *MSANTD1* or *RGS12* upstream from *HTT*, enabling missplicing from these transcripts into *HTT* exon 1. To test this notion, we re-aligned RNA-seq data to mm39 plus two tandem copies of the integrated region (8kb-*HTT*-60kb, Supp. Note C). This revealed chimeric *RGS12*-*extendedHTT* transcripts resulting from the tandem BAC insertion at levels similar to *extendedHTT* in humans (~2.3% as abundant by sequencing depth, and ~12.7% by splice junction counts, relative to *HTT* exons 2-6; Supp. Note C). The chimeric transcripts initiate within *RGS12* and are spliced to the *extendedHTT* exons integrated with the BAC array (Supp. Note C). Chimeric *RGS12* and *RGS12*-*extendedHTT* transcripts were also spliced to *HTT* exon 2 (Supp. Note C), as observed in LCLs and brain tissue. These fusion transcripts are poised to undergo repeatlength–dependent mis-splicing to *HTT* exon 1, potentially creating novel AUG-initiated ORFs encoding polyserine and polyalanine proteins, analogous to *extendedHTT* in humans. Existing RNA-seq data for the BAC-CAG model covers the ages of 2, 6, and 12 months^2^. Although we did not detect mis-splicing to exon 1 in existing RNA-seq datasets from 2-, 6-, and 12-month-old BAC-CAG mice (Supp. Note C), this is consistent with prior reports that polyserine proteins were detectable only after 18 months^2^.

The second model where aberrant translation has been observed is the N171-82Q mouse^3^. This mouse expresses an N-terminal *HTT* fragment with 82 glutamines under control of a promoter derived from the mouse major prion protein (*Prnp*)^4^. This transgene includes not only the minimal *Prnp* promoter, but also exon 1, intron 1, and part of exon 2 of Prnp^4^ (Supp. Note D). Introns are often incorporated to enhance nuclear export of transcripts, but expanded CAG repeats cause length-dependent mis-splicing, as described by us and others^5–8^. Such events could generate unexpected transcripts bearing novel AUG-initiated ORFs. Furthermore, the N171-82Q mouse has >10 insertions of the transgene^4^, raising the possibility that one or more integrated into host genes (Supp. Note D). The resulting chimeric transcripts could then undergo mis-processing to produce AUG-initiated ORFs. Other integration events (e.g., transgene fusions, inversions, or concatemers) are also possible^9^, further increasing the likelihood of aberrant RNA processing.

**Supplemental Note.**
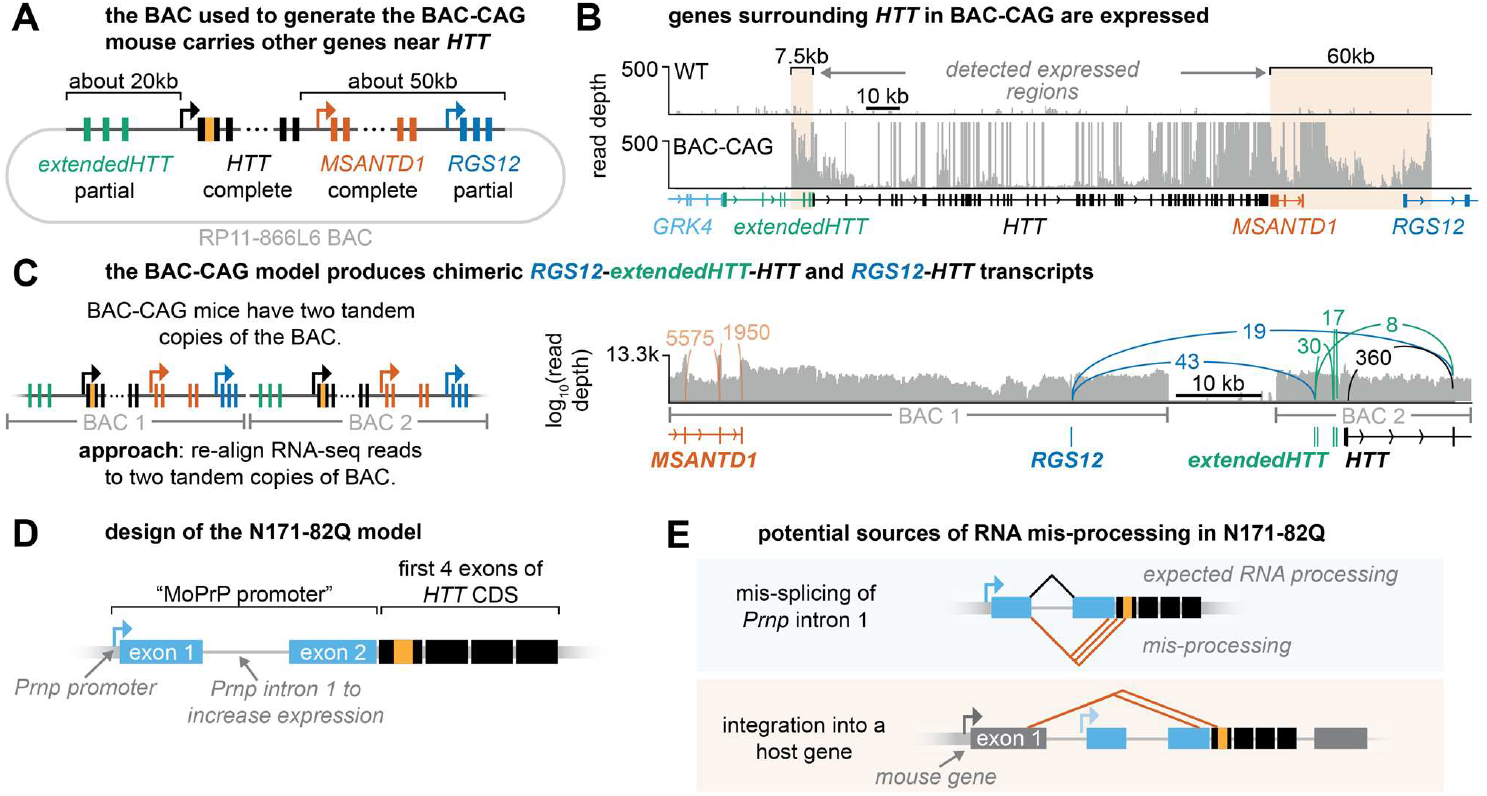
RNA mis-processing in mouse models of RAN translation. **A**. Schematic for the RP11-886L6 BAC, based on the regions reported to be present on the BAC by Gray et al. **B**. RNA-sequencing data generated from the BAC-CAG mouse at 2, 6, and 12-months were aligned to the mouse genome and human *HTT* region, then pooled to determine which regions of the BAC were integrated and expressed. We detected expression from regions approximately 7.5kb upstream and 60kb downstream from the *HTT* gene. **C**. The BAC-CAG model has been described to carry two tandem copies of the BAC. Thus, we re-aligned to the mouse genome and two tandem copies of the detected regions (as in panel B). Here, we found expression of *RGS12* fusion transcripts, where *RGS12* exon 1 was joined to the last four exons of *extendedHTT* or to *HTT* exon 2. **D**. Schematic for the transgene present in the N171-82Q model, showing that the MoPrP promoter used in this transgene includes an exon upstream from the HTT cDNA. **E**. The N171-82Q mouse model may also display mis-splicing if the expanded CAG repeat disrupts processing of MoPrP intron 1, or if the transgene is inserted into a host gene. This mis-splicing could allow foreign sequences to be joined to *HTT* exon 1, potentially generating AUG-initiated polyalanine and polyserine proteins.

Taken together, these observations suggest that aberrant translation in mouse models lacking the endogenous *extendedHTT* promoter may nonetheless arise through similar mechanisms involving mis-processing of unexpected transcripts. These context-dependent RNA processing events provide a straightforward explanation for why only some models produce polyalanine and polyserine proteins.

---

**Supplemental Figure 1.**
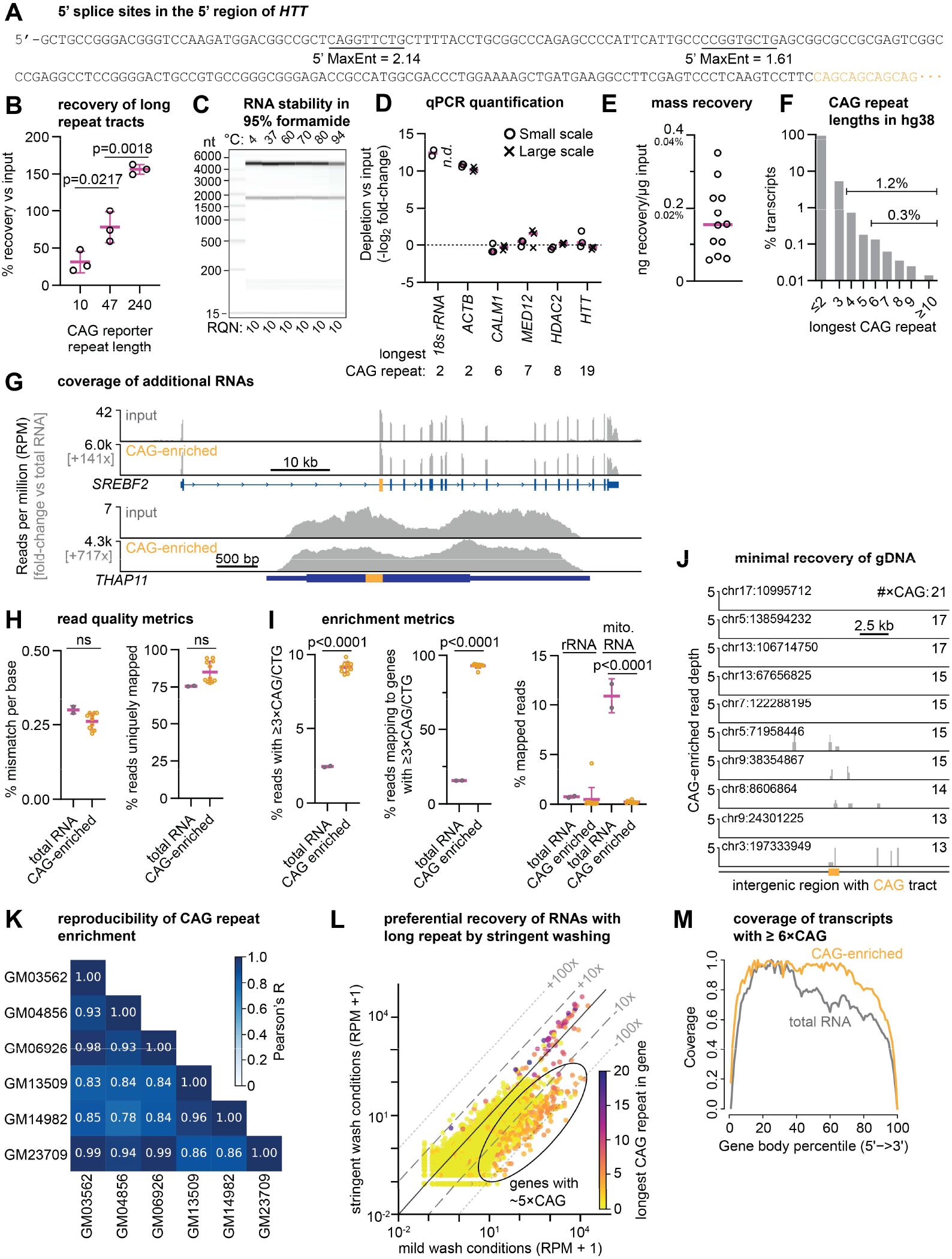
Affinity purification of tandem repeats enriches full-length RNAs with repetitive regions. **A**. Sequence of the 5’ region of *HTT* and potential donors, showing the *HTT* transcript lacks strong 5’ splice sites upstream from the CAG repeat. **B**. Real-time quantitative PCR quantification of the recovery of CAG repeat RNA reporter^13^ RNAs relative to input, after pulldown and elution with 95% formamide. Data show the mean ± SD for three independent RNA pulldowns. Recovery >100% might result from disruption of RNA structure by formamide treatment. **C**. Virtual gel from Bioanalyzer following treatment of RNA with formamide. Total RNA was incubated in 95% formamide elution buffer (95% formamide, 10 mM EDTA, and 10 mM tris pH 8) for 10 minutes at the indicated temperatures, then diluted and run without clean up. The RNA quality number (RQN) is unchanged, indicating no change in RNA integrity. **D**. Quantitative PCR analysis of several genes following small scale (10 μg) and large scale (≥ 500 μg) affinity purifications from total RNA derived from various LCL cell lines. Genes without long CAG repeats (18s ribosomal RNA, *ACTB*) are greatly depleted, while genes with repeats are largely unchanged. Each dot represents an independent RNA pulldown, with the median indicated as a purple line. **E**. Recovered masses normalized to input mass for large-scale purifications, as measured by Nanodrop. **F**. Analysis of the longest pure CAG repeat present in each transcript annotated in NCBI RefSeq for hg38. Most transcripts have no more than two tandem CAG repeats in their exons. **G**. Additional representative genes showing enrichment of CAG-containing genes, as in Fig. 1B. **H**. Read quality metrics for bulk and CAG-enriched RNA-seq. *Left:* mismatch rate, per base, as calculated by the STAR aligner. *Right:* % of reads uniquely mapping, a measure of read quality, as calculated by the STAR aligner. Each dot is one library, summarized as mean ± SD. **I**. Broad enrichment metrics. *Left:* % of reads exactly containing three or more tandem CAG/CTG repeats, quantified by bbtools. *Center:* % of reads mapping to a gene with three or more exonic CAG repeats, quantified by featureCounts. *Right:* % of mapped reads mapping to ribosomal or mitochondrial RNAs. Both types of RNA have no repeats larger than three tandem CAGs. Each dot is one library, summarized as mean ± SD. **J**. Read coverage on intergenic CAG repeats for the GM13509 library, related to Fig. 1E. **K**. Pearson’s correlation coefficient for featureCounts output (per-gene read counts, RPM normalized) for genes with ≥ 6×CAG repeats across six LCL cell line RNA pulldowns performed independently. **L**. Per-gene enrichment of RNAs with CAG repeats showing de-enrichment of RNAs with short repeat tracts using stringent washing conditions (low salt and 25% formamide; see Methods). Each dot represents one gene, colored to reflect the longest exonic CAG repeat in the gene. **M**. Relative coverage of transcripts with ≥ 6×CAG after short-read sequencing, using the RSeQC geneBody_coverage package. This shows that intact RNAs are recovered.

**Supplemental Figure 2.**
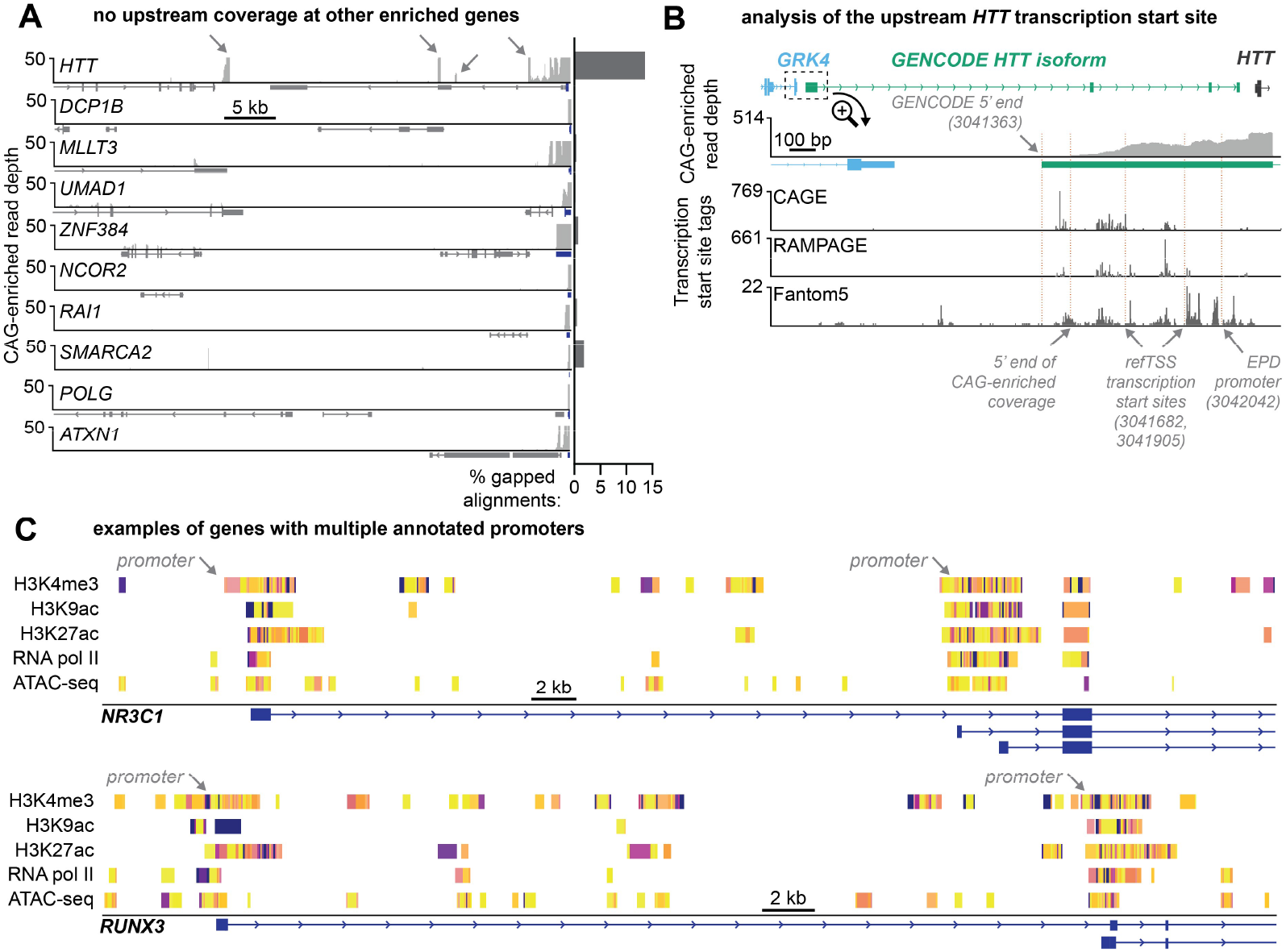
Additional evidence for the upstream HTT promoter. **A**. *Left:* CAG-enriched RNA-seq for regions 50-kb upstream from other genes with CAG repeats, showing that read density is not increased except at *HTT*. The listed gene is shown with a blue gene model, and other genes in the region (if any) are indicated by a grey gene model. *Right:* % of reads mapped in the 50-kb region having gapped alignments, which are suggestive of RNA splicing. **B**. Comparison of transcription start sites tags, annotated transcription start sites, and annotated promoters in the *extendedHTT* region. **C**. Related to Fig. 2D, additional examples of genes with two or more annotated promoters, which appear similar to the *HTT* region.

**Supplemental Figure 3.**
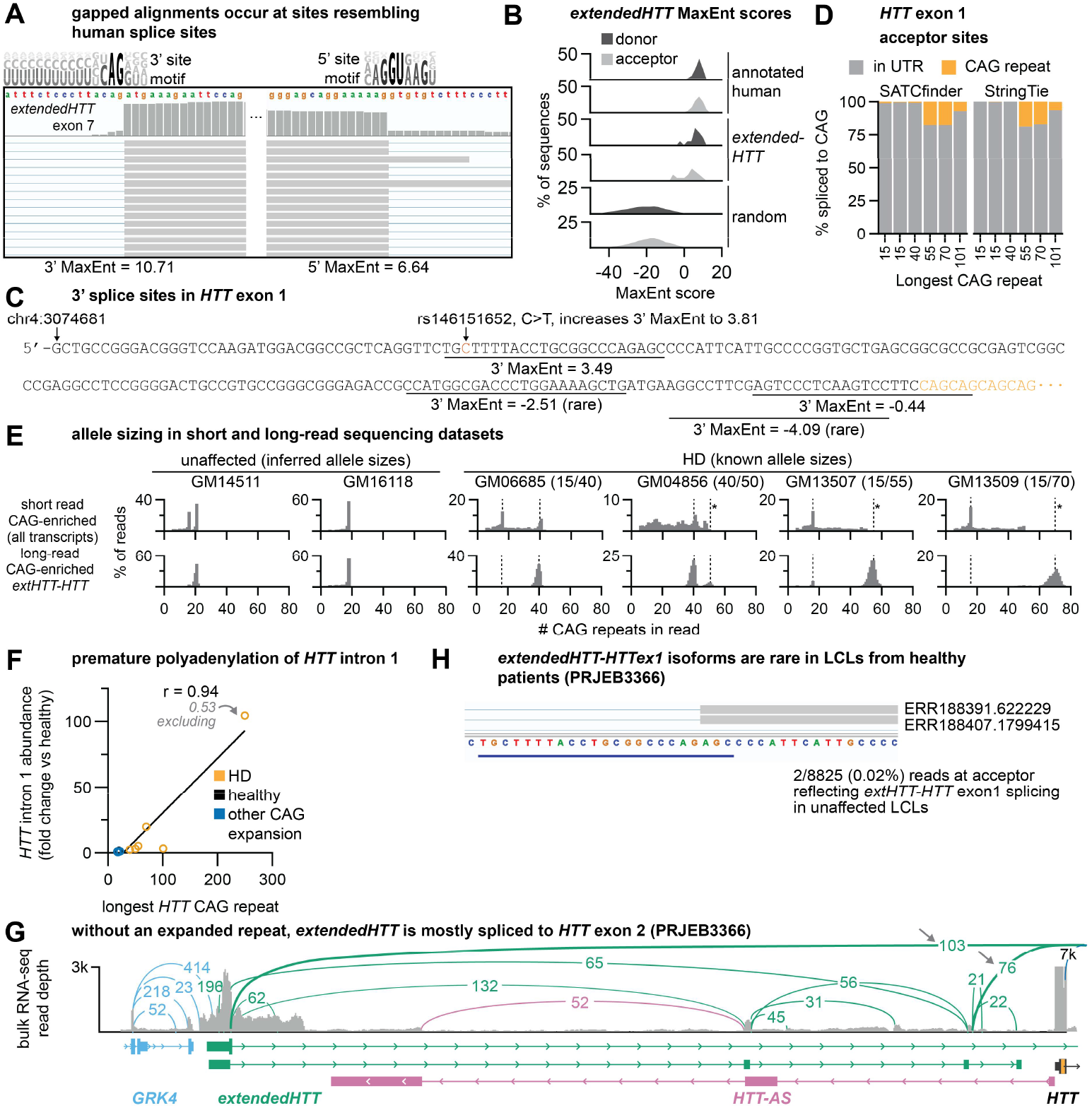
Analysis of the extendedHTT transcript. **A**. Example IGV trace for an exon of *extendedHTT*. The human consensus 5’ and 3’ splice site motifs are shown above, which generally agree with these sites. See also Supplemental Table 1. **B**. 5’ and 3’ MaxEnt scores for all ~220k human exons, the detected exons in *extendedHTT*, and 250k randomly generated 9- or 23-mers. **C**. Analysis of 3’ splice sites detected in *HTT* exon 1. **D**. Quantification of 3’ splice site selection, related to Fig. 3B. Reads with tandem repeats often display a distribution of repeat lengths (see panel E) that complicates detection of the CAG repeat as the acceptor. Thus, to quantify isoforms reflecting splicing to CAG, we applied two orthogonal methods. *SATCFinder*: long-reads were first pre-processed to trim repetitive regions (as we previously described^12^), then aligned to the *HTT* region with minimap2. Reads with repeats that mapped far upstream were counted as being spliced to the repeat tract. *StringTie*: reads were first aligned to the *HTT* region, then post-processed to have a fixed repeat length using a custom python script. Finally, we counted the number of reads that reflecting splicing to the repeat with StringTie. **E**. Quantification of *HTT* CAG repeats from short- and long-read sequencing. We benchmarked these methods against HD LCLs with known allele sizes (dashed lines). *: 150 bp short reads cannot span expanded repeats >50×CAG in one read. **F**. Quantification of CAG-enriched read coverage on the first 100 bases of *HTT* intron 1. Consistent with prior reports^36,53^, LCL cell lines with expanded repeats show increased retention of intron 1, which is attributed to premature polyadenylation. **G**. Sashimi plot for bulk RNA-seq performed on LCLs from the 1000 Genomes Project. In the absence of an expanded CAG repeat, *extendedHTT* is primarily joined to *HTT* exon 2. **H**. Rare read traces where *extendedHTT* is mis-spliced to *HTT* exon 1 in LCLs from the 1000 Genomes Project, demonstrating that mis-splicing can occur without an expanded repeat, albeit rarely.

**Supplemental Figure 4.**
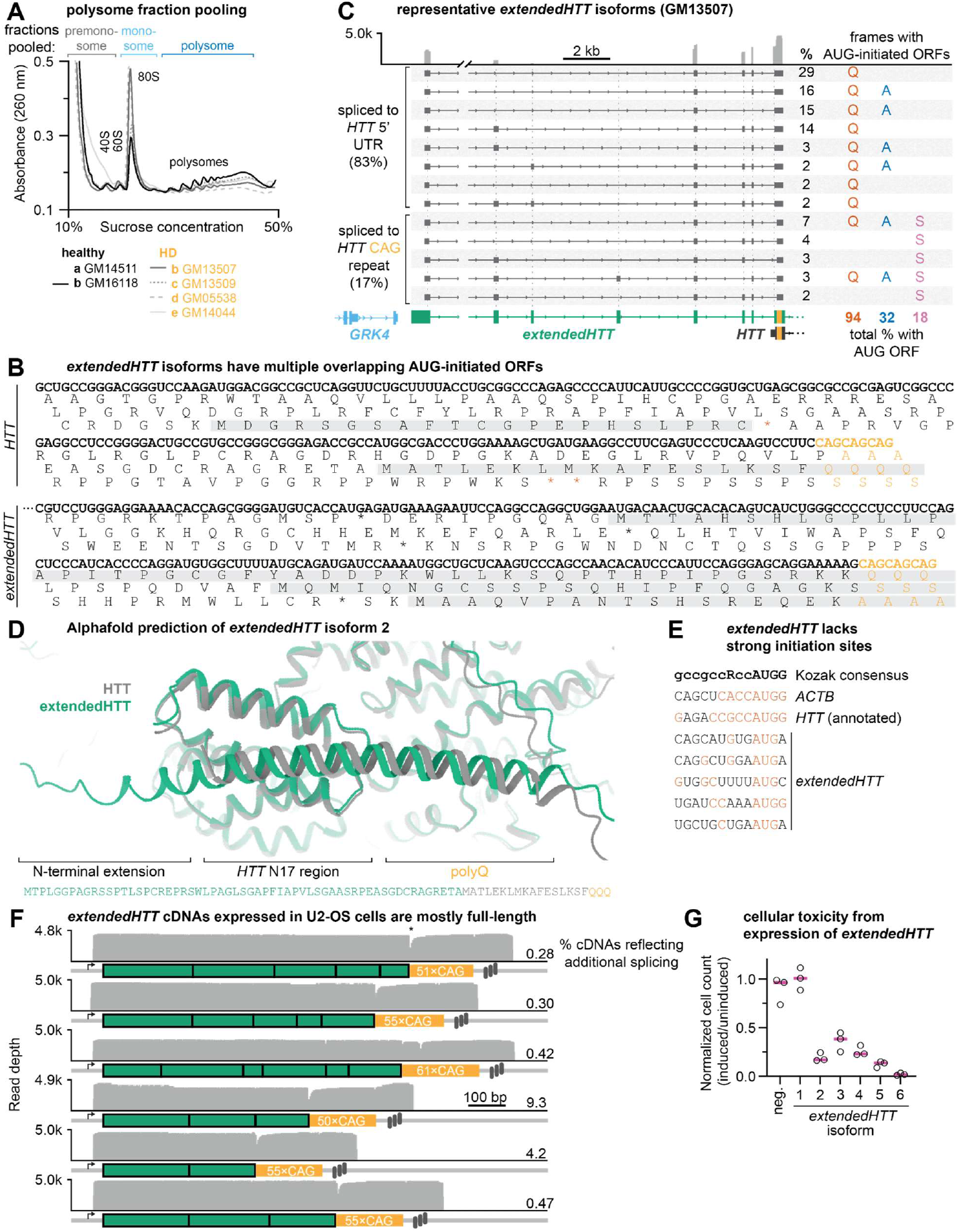
ExtendedHTT isoforms can harbor AUG-initiated ORFs across the repeat after splicing to HTT exon 1. **A**. Related to Fig. 4A, polysome profile of unaffected and HD LCLs, showing fractions pooled. **B**. Examples of open reading frames (ORFs) in the canonical *HTT* transcript and in *extendedHTT* isoform 6. Splicing to the CAG repeat shifts its sequence context, enabling AUG-initiated translation in all three reading frames. **C**. Representative isoforms for GM13507 (HD LCL) with ≥ 1% abundance from long-read sequencing. Isoform abundance and AUG-initiated ORFs that span the repeat for each isoform. Most transcripts (94%) harbor AUG-initiated polyglutamine ORFs, whereas 32% and 18% have polyalanine and polyserine ORFs. About half of *extendedHTT* transcripts have overlapping ORFs. **D**. Structures of the 5’ end of *HTT* and *extendedHTT* isoform 2 were predicted with Alphafold and aligned in ChimeraX. For this isoform, the extension is not predicted to significantly disrupt the fold of HTT (RMSD = 0.499 angstroms). **E**. Sequence alignment for *extendedHTT* AUGs as compared to the Kozak consensus sequence, where uppercase nucleotides are more important (R = A/G). *ExtendedHTT* lacks strong alignment to the Kozak consensus sequence, suggesting that leaky scanning may occur. **F**. Nanopore long-read sequencing of RT-PCR amplicons prepared from U-2OS cells expressing *extendedHTT* cDNAs, showing that the observed translation products do not result from unexpected splice variants. *: PCR-induced truncations of the repeat. **G**. Quantification of cell death from expression of *extendedHTT* isoforms for 5 days. Each data point is a separate biological replicate. Data were normalized first to the uninduced condition, then to the mock parent cell line without a repeat-containing construct.

**Supplemental Figure 5.**
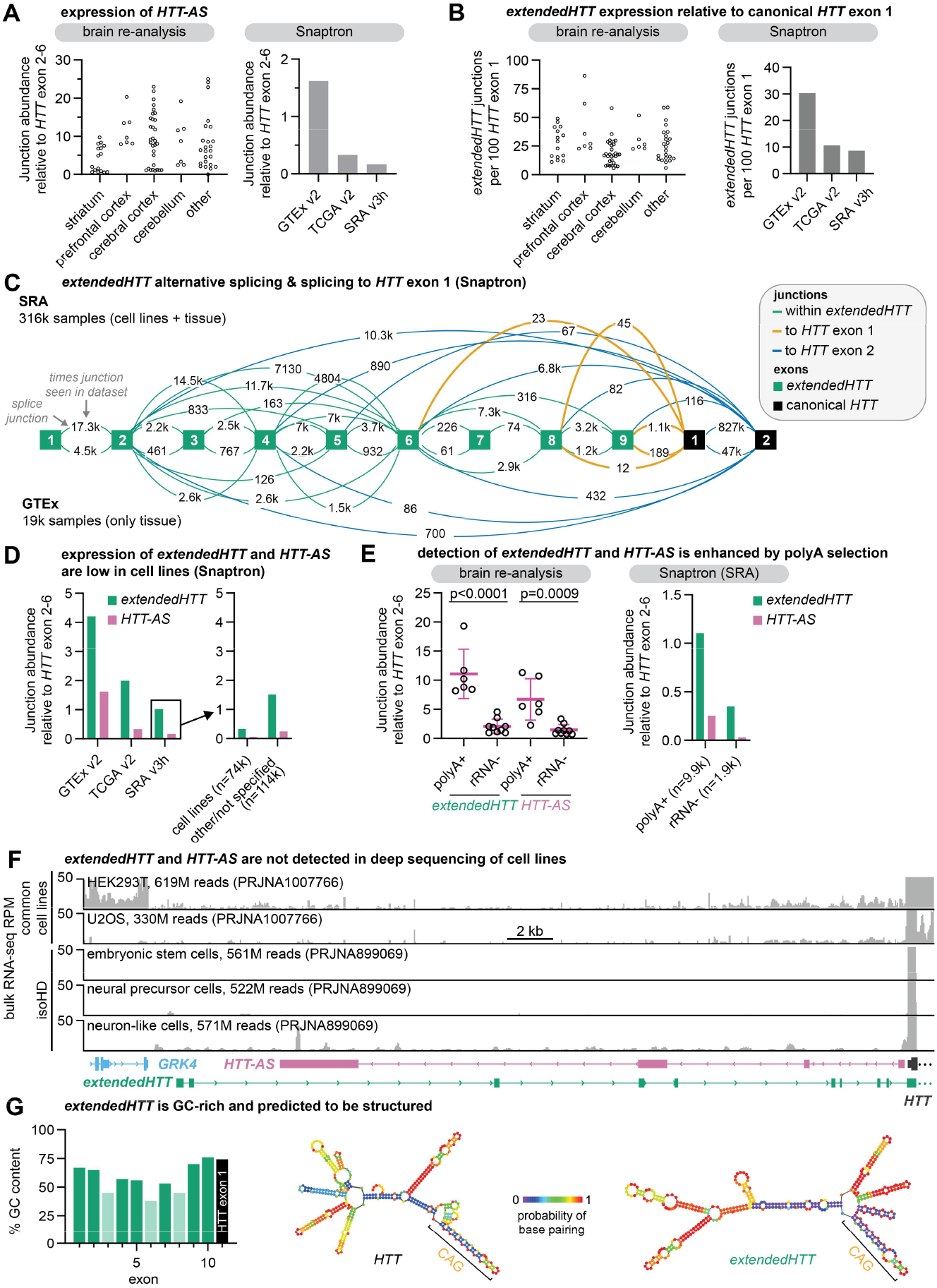
Additional evidence for extendedHTT expression. **A**. Expression of the antisense *HTT* transcript (*HTT-AS*) in brain tissues, in the re-analyzed brain dataset and in the Snaptron database of splice junctions, normalized to *HTT* exons 2-6. **B**. Expression of *extendedHTT*, as in Fig. 5B, but normalized to *HTT* exon 1. **C**. Connectivity map for *extendedHTT* and *HTT* exons in the Snaptron compilations for the NCBI SRA (Sequence Read Archive and GTEx (Gene-Tissue Expression). These results recapitulate the sequencing results in Figure 3, showing that *extendedHTT* exons are detected across numerous cell lines and tissue types. Rare mis-splicing to *HTT* exon 1 is also observed in these datasets. **D**. Quantification of *extendedHTT* and *HTT-AS* abundance relative to *HTT* exons 2-6 in the Snaptron database, related to panel C. Unexpectedly, expression of these transcripts is ~5-fold lower in samples with “cell line” in their description, with other samples (e.g. from primary cells, biopsies, or autopsies) showing higher expression. **E**. The relative abundance of *extendedHTT* and *HTT-AS* are higher in RNA-seq experiments using library preparation protocols that capture polyadenylated RNAs than in those that selectively remove ribosomal RNA (e.g., by tiling ribosomal RNA with oligonucleotides and treating with RNase H). Data are summarized as mean ± SD and significance values were calculated using Student’s t-test. **F**. Aggregated RNA-seq coverage across high-depth sequencing datasets of cell lines, similar to Fig. 5A, showing that both *extendedHTT* and *HTT-AS* are not detected in these samples. **G**. *Left*: Per-exon GC content of *extendedHTT* and *HTT* (exon 1 only). Lightly shaded exons are infrequently included on transcripts. *Right*: Minimum free energy predicted structures of *HTT* exon 1 and *extendedHTT* exon 10 spliced to *HTT* exon 1 (both with an unexpanded CAG repeat), from RNAFold. Altogether, *extendedHTT* is high GC and predicted to be highly structured.

